# Ultra Low Power, Event-Driven Data Compression of Multi-Unit Activity

**DOI:** 10.1101/2022.11.24.517853

**Authors:** Oscar W. Savolainen, Zheng Zhang, Timothy G. Constandinou

**Affiliations:** Imperial College London

## Abstract

Recent years have demonstrated the feasibility of using intracortical Brain-Machine Interfaces (iBMIs), by decoding thoughts, for communication and cursor control tasks. iBMIs are increasingly becoming wireless due to the risks of infection and mechanical failure associated with percutaneous connections. However, wireless communication increases the power consumption, and the total power dissipation is strictly limited due to safety heating limits of cortical tissue. Since wireless power is proportional to the communication bandwidth, the output Bit Rate (BR) must be minimised. Whilst most iBMIs utilise Multi-Unit activity (MUA), i.e. spike events, and this in itself significantly reduces the output BR (compared to raw data), it still limits the scalability (number of channels) that can be achieved. As such, additional compression for MUA signals is essential for fully-implantable, high-information-bandwidth systems. To meet this need, this work proposes various hardware-efficient, ultra-low power MUA compression schemes. They are investigated in terms of their BRs and hardware requirements as a function of various on-implant conditions such as MUA Binning Period (BP) and number of channels. It was found that for BPs ≤10 ms, the Delta Event-Driven method had the lowest total dynamic power and reduced the BR by almost an order of magnitude relative to classical methods (e.g. to approx. 151 bps/channel for a BP of 1 ms and 1000 channels on-implant.). However, at larger BPs the Windowing method performed best (e.g. approx. 29 bps/channel for a BP of 50 ms, independent of channel count). As such, this work can guide the choice of MUA data compression scheme for BMI applications, where the BR can be significantly reduced in hardware efficient ways. This enables the next generation of wireless iBMIs, with small implant sizes, high channel counts, low-power, and small hardware footprint. All code and results have been made publicly available.

## 1 Introduction

### 1.1 Wireless Intracortical Brain-Machine Interfaces

Brain Machine Interfaces (BMIs) are electronic devices that measure neural activity, extract informative features from that activity, and convert those features into outputs that replace, restore, enhance, supplement, or improve human functions. They can be used to treat paraplegia, quadriplegia, movement disorders, Locked-in syndrome and more [1, 2]. Intracortical BMIs (iBMIs) are the most invasive form of BMI, where electrodes are placed into brain tissue [3]. They also provide the highest resolution of BMI data, capable of measuring the firing times of individual neurons in the electrodes’ vicinity.

For iBMIs, wireless communication and powering is becoming increasingly essential, particularly for translational efforts [3–5]. Wireless iBMIs (WI-BMI) consist of a small implant in the brain that communicates wirelessly with an external decoder, where more computationally demanding processing and/or decoding is performed. The nature of the wireless channel can be either acoustic with ultrasound [6], or electromagnetic with Radio Frequency (RF) waves, where the latter is more common [4, 7–12]. The benefit of WI-BMIs is that they eliminate physical percutaneous connections (wires breaching the skin) and the associated risks (infection, mechanical damage, etc), while significantly improving the quality of life of the user. They also enable chronic powering without having to replace the implant, as battery life is limited.

The move to WI-BMIs is required for the effective translation of BMIs into clinical applications. It is one of the remaining bottlenecks. Furthermore, there is an additional desire to minimize the invasiveness of WI-BMIs by reducing the size of the implant to the mm or sub-mm scale. This is so as to minimize damage and foreign body response in the brain [3]. Ultra-minimized probe sizes can also enable ultra-minimally invasive delivery methods such as laparoscopy or injection [13]. A distributed network of free-floating, sub-mm scale implants would be ideal, because this reduces invasiveness while also improving the robustness of the system. For example, if one node failed, the BMI as a whole would continue to function. This would reduce the requirement for invasive surgery and save the user from the negative effects of their BMI failing.

As such, in terms of minimizing invasiveness and robustness, the ideal would be a system of mm or sub-mm scale free-floating, fully integrated neural recording microsystems. However, the capacity of the communication channel becomes an immediate issue. This is due to both the direct effect of data rates, but also because of heating. The same heating and channel capacity constraint holds true for monolithic WI-BMI implants as well, where one large implant measures all of the neural data from multiple channels and communicates it out along one channel.

### 1.2 Channel capacity

There is no consensus on an upper limit for channel capacity in WI-BMIs (see Table 1 of [14] for a review). However, the channel capacity is finite given neural tissue’s finite ability to absorb either acoustic or electromagnetic waves without damage. As such, the BR of each node limits the size of a distributed network of free-floating WI-BMI nodes. For example, in the case of distributed nodes communicating via a Time-Division Multiple Access protocol, the amount of available nodes is limited by the size of each node’s data packet and the channel capacity [15]. As such, if the data packet size could be reduced, more nodes could be fit into the network without risking packet collisions. For example, the Neurograins network in [15] can be scaled to 1000 channels at the maximum, but with smaller data packets, a larger number of nodes could be accommodated.

**Table 1:** Huffman encoder example [14]. For the given frequencies and codeword lengths, the compressed data has an average length of 0.8 *×* 1 + 0.1 *×* 2 + 0.07 *×* 3 + 0.03 *×* 3 = 1.4 bits, versus the 2 bits that are normally required to represent 4 symbols with a fixed-length encoding.

| Huffman Encoder |  |  |
| --- | --- | --- |
| Symbol / Key | Frequency | Codeword |
| 0 | 0.8 | 0 |
| 1 | 0.1 | 10 |
| 2 | 0.07 | 110 |
| 3 | 0.03 | 111 |

Monolithic implants suffer from the same problem: a finite channel can only communicate out so much data. Eventually, if more information is to be extracted from the brain, bandwidth becomes a constraint and on-node data compression is required.

### 1.3 Heating in WI-BMIs

Heating of cortical tissue is also major issue with WI-BMIs. This is because heating tissue can cause irreparable damage [16]. However, the acceptable heat limit in cortical tissue is not robustly understood [14, 16]. Additionally, any power that is used on-implant needs to be transferred wirelessly to the implant, some large portion of which is absorbed by the neural medium, heating it [16]. Therefore, on-implant power should be strictly limited. A 10 mW/cm^2^ heat flux is perhaps a conservative upper safety limit for cortical tissue [14], when accounting for wireless power transfer effects. For example, a retinal implant device was found to provide a heat flux of 15.5 mW/cm^2^ when including the size of the insulation, and the temperature increases were less than a degree [16, 17], which is assumed to be acceptable. However, in the same study, when the power was increased to represent 62 mW/cm^2^, the temperature increases reached as high as 3°C [16, 17], which is unacceptable. There is no equivalent study in neural tissue, but it warrants erring on the side of caution. Therefore for now we work with a 10 mW/cm^2^ heat flux.

For an example 2.5 mm *×* 2.5 mm scale, freely-floating wireless integrated neural recording microsystem, this gives an upper power budget of 1.25 mW, assuming equal heat flux from both faces and negligible heat flux from the sides of the implant. If the entirety of the available implant power budget goes into communication and one has a state-of-the-art FPGA-based RF-backscatter WI-BMI communication energy of 20 nJ/bit [14, 18], this produces a maximum BR of 62.5 kbps. The raw broadband signal bandwidth, which is typically sampled at some 16 bits/sample at 20 kHz, has a Bit Rate (BR) of approximately 320 kbps/channel. As such, the bandwidth would need to be compressed by 80% to allow even a single broadband channel to be communicated off-implant. In practice, reported communication energies are typically not measured *in* or *ex vivo*, and the communication energy of the same system in the neural medium would be significantly higher. For example, the lowest FPGA communication energy that has been shown was 20 nJ/bit [14, 18], but this involved a head-mounted wireless communication board external to the cranium of the free-moving rat subject. As such, with practically higher communication energies, the number of channels that can be fit on-implant is fewer.

Additionally, one has to account for the hardware static power, where even ultra-low power FPGAs like the Lattice ice40LP1K require over 100*μ*W for the static power consumption. Furthermore, we need to account for the front-end pre-amplifier and ADC power costs [14]. Therefore, if we are going to communicate anything off-implant in mm-scale free-floating WI-BMIs, some form of data compression will be required.

### 1.4 Data Compression in WI-BMIs

#### 1.4.1 Lossless Compression

Data compression comes in two forms: lossless and lossy. Lossless compression consists of giving shorter codewords to more common signals, which is the basis of Information Theory [19]. There are many lossless compression algorithms, however most of them require too many resources to be practical for extremely hardware-constrained devices like WI-BMIs, e.g. Lempel-Ziv, Arithmetic, or Adaptive Huffman encoding. A lossless compression algorithm that works well for WI-BMIs is Static Huffman (SH) encoding, since it can be implemented using only a small number of Look-Up Tables (LUT). It works by one pre-training a Huffman encoder [20] on some representative data, and then implementing the encoder on-implant with LUTs. If the data that is measured on-implant is well-represented by the training data, the encoder will efficiently compress the data. An example of a Huffman encoder is shown in Table 1. SH encoders were used to great effect in [14] to compress Multi-Unit Activity (MUA) signals. They were also used in [21] to compress other intracortical signals, e.g. Entire Spiking Activity (ESA), Local Field Potential (LFP) and Extracellular Action Potentials (EAP). Huffman encoders are optimal among methods that encode symbols separately. Given their hardware efficiency, in their static form, they are attractive for WI-BMIs.

#### 1.4.2 Lossy compression

The second form of data compression is lossy compression. One eliminates information that is assumed to not be of interest to the final application, reducing the bandwidth. This is also called feature extraction, dimensionality reduction, etc. In experimental neuroscience, it is standard to do lossy compression for neural signals via signal processing techniques. A typical data flow is where the raw signal is processed with Principal Component Analysis (PCA) or wavelets, the resulting features are clustered to perform spike-sorting, and this outputs Single-Unit Activity (SUA) spike trains which are analyzed. This reduces the BR while improving the interpretability of the data.

However, the above procedure uses what are called ‘data adaptive’ methods. Data adaptive methods are powerful in that they can use statistics within the signal to tailor their processing to the signal. Common examples in neuroscience are PCA, ICA and t-SNE. However, they typically consume significant computing resources. As such, on-implant in BMIs, it is typical to do lossy compression via ‘data agnostic’ methods. These are methods where the processing procedure is a fixed pipeline, where the same computations occur regardless of the incoming signal. There are a few data agnostic features, so to speak, that are commonly extracted in iBMIs. These, along with their typical BRs, are shown in Fig. 1. These can be divided into two categories: LFP features and EAP features. LFPs consist of the lowpassed broadband at 100-300 Hz, and are believed to result from the sum of extracellular currents and spike activity in the vicinity of the electrode [22, 23]. While low bandwidth, easy to measure and chronically available, it has not been shown to have as good decoding performance as higher-frequency features such as those derived from the EAP [24].

**Figure 1:**
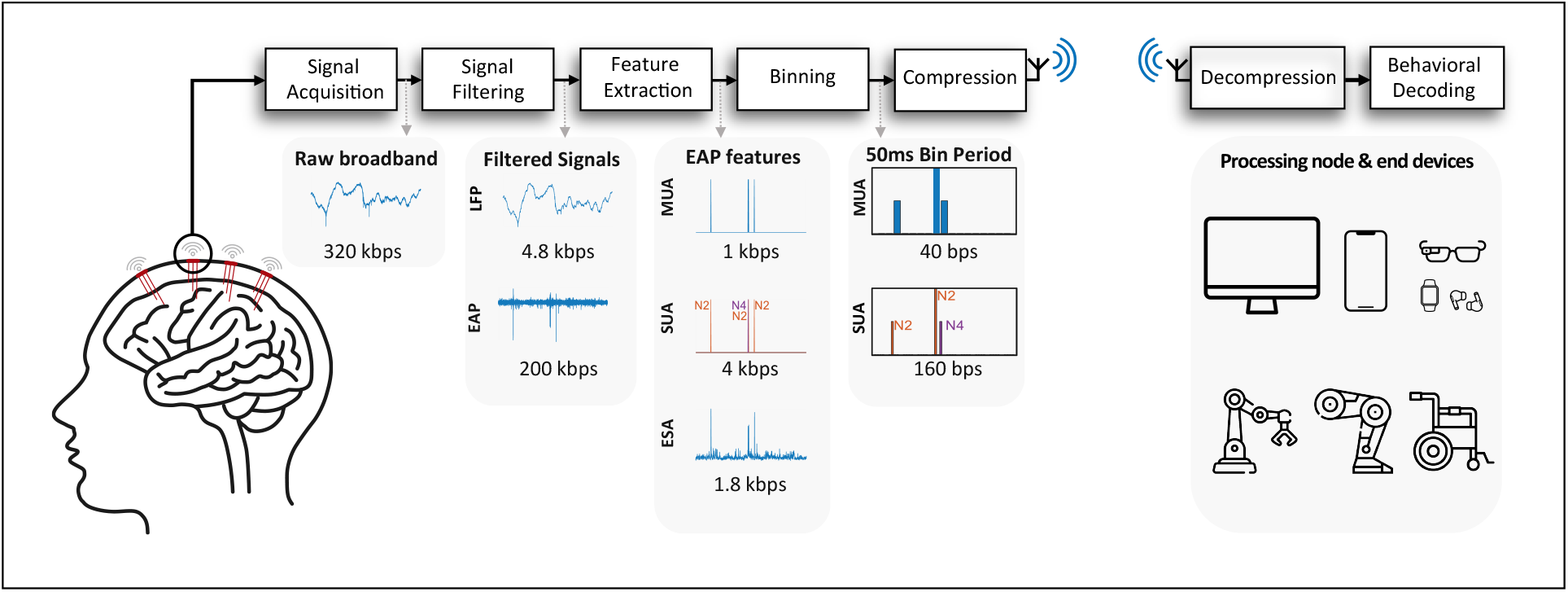
Reprinted from [14] with permission. Typical BMI data processing and compression flow, with common extracted features / lossy compressions of intracortical broadband data. The numerical values beneath the signals give approximate BRs per channel for that signal. LFP: Local Field Potential, equal to the lowpassed broadband at approx. 300 Hz. EAP: Extracellular Action Potential, equal to the highpassed broadband at approx. 300 Hz. MUA: Multi-Unit Activity, equal to the unsorted thresholded spike activity from the EAP. SUA: Single-Unit Activity, equal to the sorted thresholded spike activity from the EAP. ESA: Entire Spiking Activity, equal to the enveloped rectified EAP signal, giving an envelope of unsorted spiking activity.

The EAP consists of the ∼300 Hz highpassed broadband. Most of what makes up the EAP signal is spikes of varying amplitude combined with thermal and electronics noise [5,25,26]. These spikes correspond to the signatures of neurons firing in the vicinity of the electrode, and can be observed in the EAP signal in Fig. 1. The EAP signal is highly sparse, given the infrequency of these spiking events which are believed to contain most of the interesting information in neural signals. To remove the uninteresting noise, it is common to extract various features from the EAP, e.g. SUA, Multi-Unit Activity (MUA) and Entire Spiking Activity (ESA).

Whereas SUA involves clustering each spike into a putative neuron based on spike shape, the MUA representation assigns each spike on the same electrode to the same putative neuron. Although evidence is somewhat mixed, it is generally believed that MUA gives very similar Behavioral Decoding Performance (BDP) to SUA [24, 27–29]. Additionally, the spike-sorting procedure is very computationally expensive to perform on-implant [30, 31], and so the MUA signal has gained significant popularity, and is now considered the default signal in modern WI-BMIs [4, 5, 32]. By reducing the broadband to only the spikes in the SUA or MUA features, significant *de facto* compression is achieved, along with power savings [5, 30]. As such, this work will focus on exploring WI-BMI data compression in the context of the MUA signal. In particular, we will analyse it in the context of mm-scale free-floating, fully integrated neural recording probes, however the results generalise to other WI-BMIs.

### 1.5 Multi-Unit Activity

As aforementioned, when a neuron fires in the vicinity of an intracortical electrode, the electrode measures a short spike in voltage. In the MUA signal, the timing of this spike is noted, and the firing times binned at some resolution, i.e. the Binning Period (BP). Therefore, at each BP, each electrode channel outputs an integer value representing the MUA Firing Rate (FR), equal to how many spikes were detected by the electrode during the BP.

The MUA BP is typically 1 ms [5], since spikes last approximately 2 ms. The dynamic range *S* is also typically set to a binary output, i.e. *S* = 2. This produces *ceil*(*log*_2_(2)) = 1 bit/sample, which at a 1 ms BP corresponds to a bit rate (BR) of 1 kbps/channel. In fact, this is technically identical to 1-bit sampling at 1 kHz. This is a significant reduction over the raw broadband signal bandwidth of approximately 320 kbps/channel. However, a 1 kbs/channel is still too high for many WI-BMI applications.

For example, for the aforementioned 2.5 mm *×* 2.5 mm scale, freely-floating wireless integrated neural recording microsystem, each node has an approximate upper power budget of 1.25 mW. With a 1 kbps/channel BR and an ultra-low communication energy estimate of 20 nJ/bit, this corresponds to 62 channels. In practice, if one accounts for the required MUA processing power, front-end pre-amplifiers and ADC, practically higher communication energies (e.g. 100 nJ/bit), and hardware static power, it is unlikely that even 10 MUA channels could be hosted on such an implant. As such, it warrants determining whether easy-to-implement, hardware-efficient data compressions schemes exist that can enable significantly higher channel counts per node, improving the coverage and performance of WI-BMIs.

### 1.6 This Work

A hardware-efficient MUA compression system was introduced in [14]. For its lossy element, it used binning of the MUA signal and thresholding the dynamic range of the resulting signal. For the binning, this is because increasing the BP is a simple way of compressing the MUA signal [5, 14, 33]. However, because the BP is the main temporal bottleneck in the whole BMI, increasing BP reduces the temporal resolution of the final decoded behavior. For example, if the BP is 100 ms, the BMI user will only be able to control the behavioral module of the BMI with 100 ms temporal precision. As such, the BP should be minimised if possible.

For the range thresholding, it was found that the dynamic range *S* of the signal can be thresholded to further compress the data [14]. This is done by taking all MUA FR values above *S* − 1, and setting them to *S* − 1, for some integer value *S* chosen by the researcher. For various BPs, it was found that *S* could be reduced significantly without negatively affecting the BDP, across multiple Non-Human Primate (NHP) subjects and datasets [14].

Shorter BPs make the firing rates sparse. In other words, in a smaller time window it is more likely that a FR of 0 will occur. Using this known property of MUA signals can help us improve the compression performance for BPs below 50ms. As such, in this paper, we reproduced the work in [14] as a baseline method. This method is referred to as the ‘windowed’ architecture, where each channel outputs its data for each BP. We also propose several event-driven architectures, where only data from active channels is transmitted for the given BP, taking advantage of sparse MUA FRs at lower BPs. Overall, four compression schemes will be examined: the windowed scheme from [14], an Explicit Event-Driven (EED) method, which transmits the active channel IDs with corresponding spike counts, a Delta Event-Driven (DED) method, which transmits the delta-sampled active channel IDs with corresponding spike counts, and a Group Event-Driven (GED) method which uses a form of run-length encoding.

### 1.7 Original Contributions of This Work

This work contains multiple original contributions to MUA compression. These are as follows:

- Design and analysis of novel MUA data compression architectures, such as the EED, DED, and GED encodings.
- The hardware designs, made publicly available, of these encodings.
- Comparison of these encodings to the state of the art in MUA compression. Very large improvements were made in terms of communication bandwidth for BPs below 20 ms, appropriate for higher temporal resolution MUA-based BMIs. Compression performance was improved by up to over an order of magnitude for smaller BPs relative to the state of the art. This allows for significantly more MUA channels to be fit on-implant while staying within heating limits. This was found to be true using data from 3 different publicly available datasets, including 5 NHP subjects and 23 hours of Utah array multi-channel recordings.
- We showed that SH encoders, trained on distributions that make simple assumptions about MUA data, perform just as well as non-causal Adaptive Huffman encoders while being far more hardware-efficient and causal. As such, SH encoders seem to be a very attractive means of data compression for MUA-based WI-BMIs.
- We investigated the use of sample histograms for mapping, as in [14], for the event-driven schemes proposed in this work. We determined their effect on BR and processing power, as well as analyzing trade-offs in their use.

The rest of this paper is organised as follows: Section 2 introduces three datasets used in this work and details the compression methods. Section 3 shows the compression performance and hardware cost of each method. Based on these results, we give the recommend compression settings at different BP and channel counts, trading off between the BR, processing power and hardware cost. Section 4 discusses some algorithm and hardware design considerations in MUA compression and Section 5 concludes this work.

## 2 Methods

All compression algorithm work was done in MATLAB R2021A. All hardware design and optimisation work was done using ModelSim Lattic Edition and IceCube2020.12. All code and results are publicly available at [34].

### 2.1 Dataset and Data Formatting

Brain signal conditions can vary across subjects and tasks. In order to reduced the bias, we have used three different publicly available datasets [35–37], summarised in Table. 2. For each dataset, the SUA data was intra-channel collated to MUA, then binned to the desired BP, where BP ∈ [1, 5, 10, 20, 50, 100] ms. Based on the results in [14], we limited *S* to [2, 2, 2, 2, 3, 5] for BPs of [1, 5, 10, 20, 50, 100] ms respectively. The Supplemental Material includes results from *S* values ∈ [ℤ^+^, 2, 3, …, 14, 15].

**Table 2:** Dataset summaries.

| Dataset | Neural data type | Species, electrode type and brain region | Details | Behaviour |
| --- | --- | --- | --- | --- |
| **Flint [35]** | SUA | Rhesus macaque monkey<br>Utah array<br>M1 | One subject<br>12 recordings across 5 days<br>96 channels<br>Recording lengths (quartiles, s):<br>597, 604, 630 | Free-reaching hand task<br>Continuous data stored |
| **Sabes [36]** | SUA | Rhesus macaque monkeys<br>Utah array<br>M1 and S1 | Subjects Indy and Loco<br>37 recordings for Indy across 10 months<br>10 recordings for Loco across a month<br>96-192 channels<br>Recording lengths (quartiles, s):<br>Indy: 472, 524, 816<br>Loco: 1771, 1928, 2384 | Free-reaching hand task<br>Continuous data stored |
| **Brochier [37]** | SUA | Rhesus macaque monkeys<br>Utah array<br>M1 and PMv/PMd | Subjects N and L<br>Single session recordings<br>96 channels<br>Recording lengths (s):<br>N: 1003<br>L: 709 | Hand reaching task<br>Target stored |

The length of the recordings was largely irrelevant for the sake of this work, as all encoding was done without using a time derivative and the bandwidth was measured in [bps/channel], normalising for time. As such, a standard length of 100 s was set for each recording, long enough to gather a stable distribution for each channel for any of the tested BPs. To maximize the use of the data in observing the effect of channel count, recordings were split into consecutive 100 s segments and collated together. For example, a 400 s recording was represented as 4 parallel channels of 100 s long recordings, where *x*_*o*_,…,*x*_*N/*4−1_ became one channel, *x*_*N/*4_,…,*x*_*N/*2−1_ became the next, etc., where *x*_*k*_ is a single-channel MUA recording and *N* = 400 *s/*BP is the length of the recording in samples. A total of 79200 channels were available after splitting and collation.

The data was then split into training and testing sets. This is because the encodings have parameters that need to be optimised, which was done on the training set. The testing of parameter-optimised encoding on the testing set can then provide an objective estimation of the encoding performance on the unseen data. The training-testing split was done by randomly selecting 30000 channels and placing them into the training set. The remaining 49200 were put into the testing set. For each encoding, BP, *S* and *n* combination, the training and testing results were each averaged across 5 training and testing runs. For each run, *n* ∈ [10, 100, 1000, 10000, 30000] channels were selected randomly without replacement from amongst all training or testing channels.

### 2.2 Windowed encoding

The windowed encoding [14] was the first scheme to be implemented. It is called “windowed” because it sends out the number of MUA events in a non-overlapping window of length BP for each channel, regardless of whether an event occurs on a channel or not. It consists of representing the multi-channel MUA data as multiplexed data. Without lossless compression, the length of the data block is *n × m*, where *n* is the number of channels and *m* is the number of bits used to represent the MUA FR per BP on each channel. An example using a standard binary representation with *m* and *n* = 3:

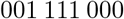

would indicate that 1 neuron fired (001) on channel 1, 7 neuron firings occurred (111) on channel 2, and 0 (000) on channel 3. An advantage is that the channel ID is implicitly encoded in bit position, and so does not need to be explicitly encoded. E.g.,

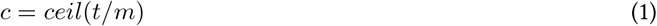

where *c* ∈ [ℤ, 1 ≤ *c* ≤ *n*] is the channel ID, *t* ∈ [ℤ, 1 ≤ *t* ≤ *n ×m*] is the bit position and *ceil* is the ceiling function. *m* is the number of bits required to represent all possible MUA FRs losslessly, and is generally set as *ceil*(*log*_2_(max(*X*) + 1)), where *X* is the multi-channel MUA data. However, as discussed in Section 1.4.2, one can lossily compress the data by limiting the dynamic range at an *S* value, setting *X*[*X >* (*S* − 1)] ← (*S* − 1). It requires

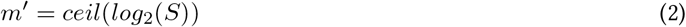

bits to represent a range of 0 to *S* − 1 MUA FRs. If *m*′ *< m* = *ceil*(*log*_2_(max(*X*) + 1)), this lossily reduces the required bandwidth. As such, the windowed encoding typically has a BR of:

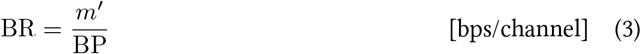

However, as in [14] we integrated the windowed encoding with a SH encoder that compressed the FRs. The SH encoder was pre-trained on a decaying exponential which mimics the distribution of MUA FRs at BPs ≤ 100 ms [14], where smaller codewords are given to smaller FRs.

### 2.3 Explicit Event-Driven encoding

In the windowed architecture, the FR of each channel is encoded. The channel ID is implicitly encoded in bit position. To the best of the authors’ knowledge, the following event-driven architectures are proposed for the first time in MUA compression. In these event-driven architectures, the channel ID is explicitly encoded and only active channels will be transmitted, i.e. the channel ID is only sent out if a non-zero FR occurs on that channel, followed by a binary codeword representing the rate. If a channel has a FR of 0, nothing gets communicated for that channel, and the offiine decoder assumes the missing channels had FRs of 0. For sparse signals, i.e. where FRs not equal to 0 are rare, this can offer reduced bandwidth over the windowed paradigm, where even FRs of 0 need to be communicated for each channel.

As the channel IDs need to be explicitly encoded, the simplest method is to give the channels a standard binary codeword of length *k*_1_, where:

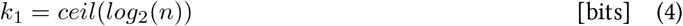

Similarly, the MUA FR for each channel is encoded as a binary codeword of length *m*_2_:

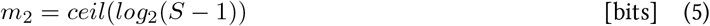

An exception occurs if *S* = 2, i.e. *m*_2_ = 1 and *i* is limited to 0 and 1. In that case the firing rate codeword is unnecessary as the decoder can assume all received channel has the firing rate of 1. An example of the encoding without lossless compression, with *n* = 4, *k*_1_ = 2 and *m*_2_ = 2, is given by:

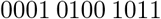

This shows that channel 1 (00) had a FR of 2 events (01) in the bin, channel 2 (01) had FR = 1 (00), channel 3 (10) had FR = 4 (11), and channel 4 had a FR = 0 (absent). In this work, this encoding was named the explicit event-driven (EED) encoding because the FR per channel is explicitly encoded in the *m*_2_-length codeword that follows the channel ID.

To include the SH encoder, there was no good way to compress the channel IDs, since they are *a priori* equally likely to be active. Therefore, the EED encoding uses the same *k*_1_ length codeword for the channel IDs, but uses varying length SH codewords for the FRs. As in the SH windowed implementation, the FR encoder was trained on a decaying exponential.

### 2.4 Delta Event-Driven encoding

The previous encoding suffers since the probability of each channel being active is *a priori* equal. As such, the channel IDs cannot be compressed as they are with SH encoders. However, the channel difference between two successive active channels has varying probability, and therefore Huffman encoding can be used. In other words, we can encode the delta-sampled channel IDs of active channels. Therefore, in this DED encoding, if a channel has a FR above 0, it is given a Δ value by subtracting its ID, *j*_*current*_, from the ID of the previous channel to have a FR above 0, *j*_*previous*_. I.e., Δ = *j*_*current*_ − *j*_*previous*_.

Therefore, the FRs were compressed with the same SH encoder as in the EED encoding. However, the Δ-sampled channel IDs were compressed using a SH encoder trained on a decaying exponential. Significant memory optimisation was also done by setting a maximum Δ-sampled SH encoder size, using a form of run-length encoding. This reduced BR slightly but significantly reduced memory requirements. The details of this optimisation are extensively detailed in Sections 3 and 4 in the Supplemental Material.

### 2.5 Group Event-Driven encoding

In the GED encoding, one uses position and stop symbols to encode the FRs. As in the EED encoding, one explicitly encodes the channel ID, but here one encodes the FR per channel implicitly in channel ID position. For example, in decimal,

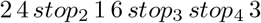

signifies that channels 2 and 4 had a FR of *i* = 1 in the given bin, channels 1 and 6 had a FR of *i* = 2, channel 3 had FR *i* = 4, and the rest of the channels had FR *i* = 0. For the codeword lengths, the simplest implementation is to give the stop symbols and the channel IDs a length of *k*_2_ bits, where:

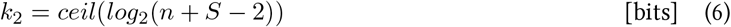

E.g. *n* = 2, *S* = 4, means that there are 2 channels, FRs between 0 and 3 inclusive can be encoded, and *k*_2_ = 2. Channel 1 gets a codeword of 00, channel 2 gets a codeword 01, and the stop symbols for *i* = 2 and *i* = 3 get codewords of 10 and 11 respectively. It essentially involves sorting the channels by FR, and using a form of run-length encoding.

There was no clear role for SH encoding, since the stop symbols and FR codewords need to come from the same encoder. Estimating these probabilities *a priori* is difficult. As such, no SH encoding for the group encoding was used.

### 2.6 Hardware Implementation

The above encoding algorithms were implemented on an ultra-low power, hardware-efficient FPGA target, Lattice ice40LP1K, in order to assess their hardware complexity (power consumption and resources usage). The results are a good approximation of the ultimate ASIC design, but enables rapid testing of algorithm parameters. The implementations are given in the Supplemental Material, Section 3.

## 3 Results

In Fig. 2, we plotted the BRs of each encoding at different BPs and *n*. We can observe that the windowed and DED encodings perform best, with the best one depending on BP and *n*. From the hardware perspective, the windowed encoding is far more hardware efficient than the DED encoding. Whilst the amount of logic cells is similar for the two encodings, the DED encoding uses significantly more memory, and therefore the processing power is higher. Additionally, the windowed scheme performs the same independent of channel count. However, the DED scheme is affected by channel count. When *n* increases, the delta-sampled channel IDs can be larger, meaning longer codewords and higher BRs. As such, increasing *n* slightly increase the DED BR, but significantly less than for the EED and GED encodings.

**Figure 2:**
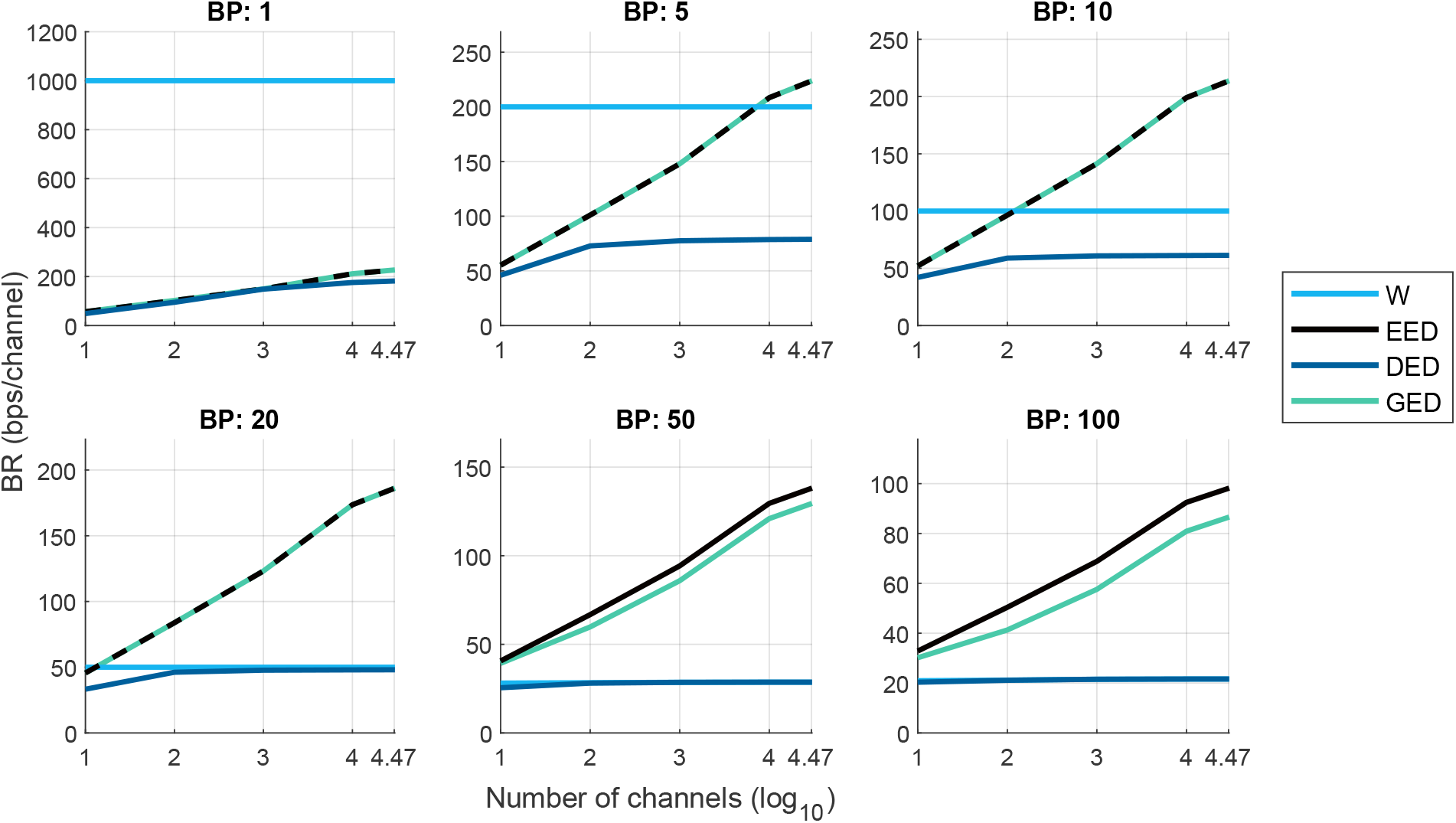
Bit Rates for communication schemes at a BP ∈ [1, 5, 10, 20, 50, 100] ms. For each BP, *S* was respectively fixed as [2, 2, 2, 2, 3, 5]. At *S* = 2, the explicit and GED encodings are mathematically identical, and so their BRs overlap for BPs ≤ 20 ms.

As such, the optimal selection generally varies as a function of channel count and BP. Ultimately, we want the total power on-implant to be reduced. As such, Fig. 3 shows the total dynamic power for the FPGA implementation, made up of the processing power for each encoding and the communication power, estimated as the BR multiplied by an estimated 20 nJ/bit communication energy.

**Figure 3:**
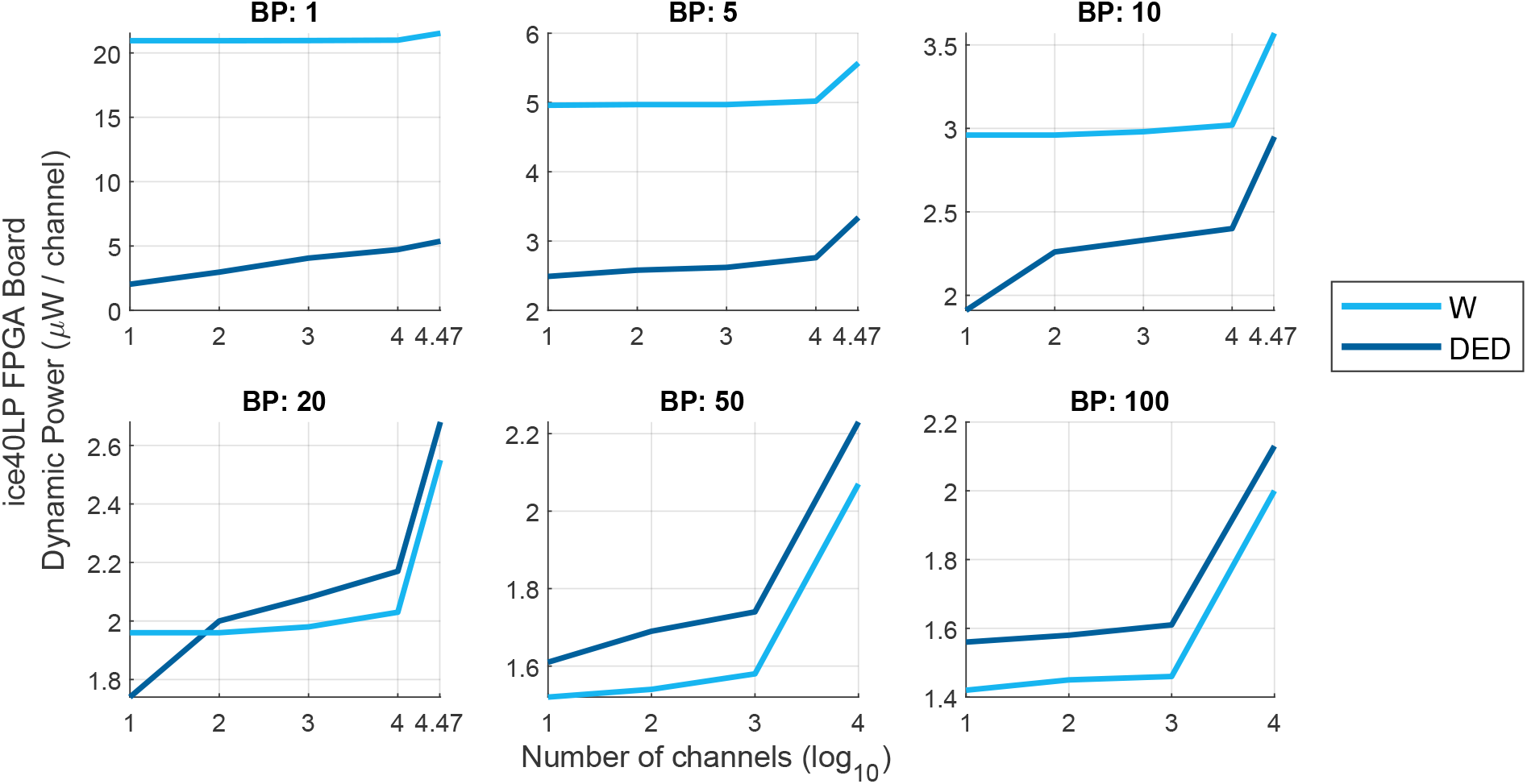
Total communication + processing power, i.e. dynamic power, for the windowed and DED compression schemes at a BP ∈ [1, 5, 10, 20, 50, 100] ms. For each BP, *S* was respectively fixed as [2, 2, 2, 2, 3, 5]. For BPs of 50 and 100 ms, the power is not given for 10000 and 30000 channels since the memory requirements exceeded the FPGA target memory budget.

### 3.1 Optimal Encoding Selection

From Fig. 3 and our analysis of the hardware costs in the Supplemental Material, for each BP and *n* we selected the optimal compression system. These are given in Table 3 (a). We then determined each selected system’s performance on the test data, i.e. data that had hereto been untouched, and the BRs are given in (b). The associated required number of FPGA logic cells and memory are given respectively in Table 3 (c) and (d). The total dynamic power on the Lattice ice40LP FPGA target, determined on the testing data, is given in (e). In summary, the DED encoding is suitable for short BPs (less than approx. 20 ms) while the windowed encoding is preferred when the channel count or BP is increased.

**Table 3:**
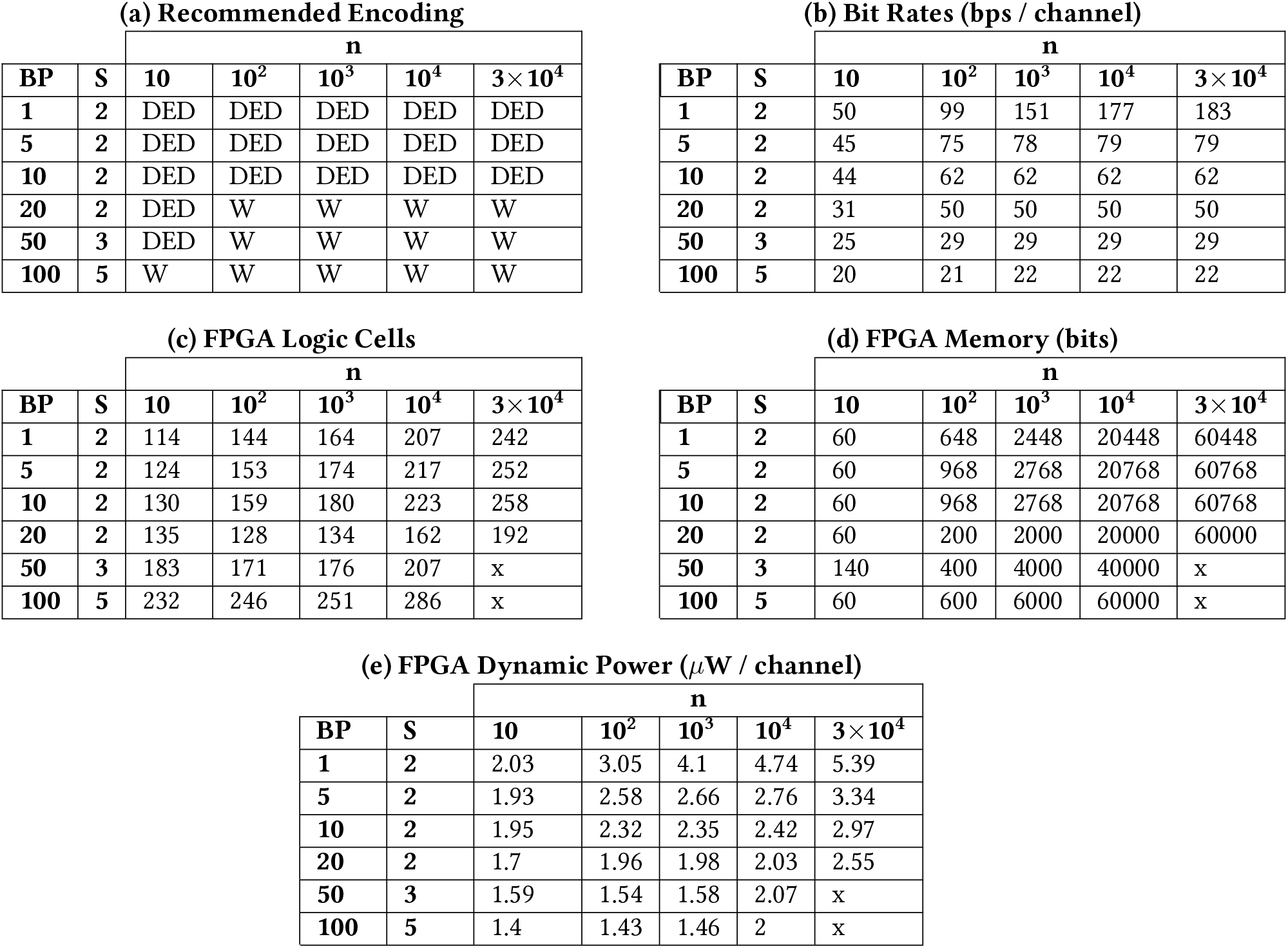
Suggested SH encodings and corresponding BRs and hardware resources for each BP and channel count. BR results from test data. ‘W’ represents the windowed encoding. ‘x’ entries are where the memory requirements exceeded those of the FPGA platform we selected. The FPGA Logic Cells and Memory are shared across all channels.

### 3.2 Testing the Encoding/Decoding

While having the BRs for each encoding and implementation is important, it is necessary to verify that each encoding functions as desired and is feasible. As such, encoding functions that encoded the binned and *S*-saturated MUA data according to each encoding and implementation, as well as with mapped versions, were implemented. The BRs were determined from the encoded data directly and were verified to match the analytically derived BRs, with some very small differences due to rounding errors. The full decodability of the data was then verified by decoding the original binned and *S*-saturated MUA data from the encodings and verifying that the original and decoded versions were a perfect match. This was done successfully for each method. The MATLAB code for each method and implementation’s encoding and decoding is given at [34].

## 4 Discussion

### 4.1 Impact of Compression on Channel Counts in FPGA Target

From this, knowing our BRs per BP and channel count, we can derive how many channels can be hosted on-implant, for different FPGA board dimensions and BPs. We assume:

- A 10 mW/cm^2^ heat flux limit;
- An FPGA static power of 162 *μ*W;
- A separate power and hardware budget for the front-end amplifiers, filters and ADCs;
- A 20 nJ/bit communication energy;
- A binner processing power of 0.96 *μ*W, independent of BP [14].

Using the information from Table 3 (e), we can derive how many channels can be hosted on-implant while staying within implant power limits. This is compared to the number of channels that can be hosted given the standard uncompressed MUA representation at 1 ms BP, *S* = 2, that has a 1000 bps/channel BR.

The results are shown graphically in Fig. 4. It can be observed that between 4.6 and 26 times more MUA channels can be fit on-implant with data compression, depending on BP ∈ *{*1, 5, 10, 20, 50, 100*}* ms and FPGA size ∈ *{*1, 2.5, 5, 7.5*}* mm. As such, one can observe that data compression allows one to fit many more MUA channels onto the same implant. Therefore, the compression schemes developed in this work are a useful addition to MUA-based WI-BMIs.

**Figure 4:**
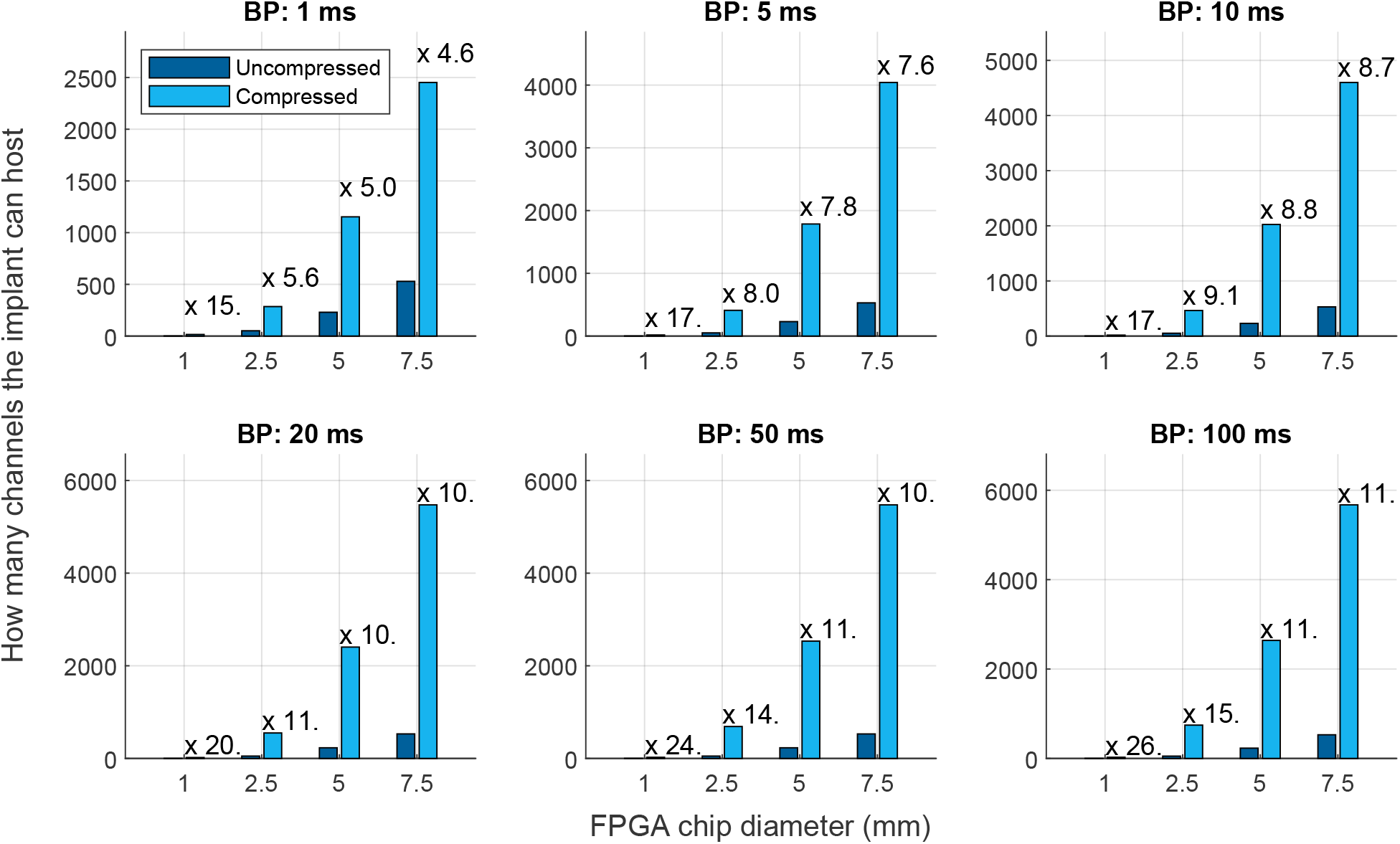
Number of MUA channels we can host on a Lattice ice40LP FPGA of different dimensions for the selected communication schemes at a BP ∈ [1, 5, 10, 20, 50, 100] ms while remaining within the power budget. This is compared to the number of channels that can be hosted given the standard uncompressed MUA representation at 1 ms BP, *S* = 2, that has a 1000 bps/channel BR. The required power for any front-end ADC and pre-amplifiers is ignored. The total dynamic power budget is given by the FPGA dimensions multiplied by a heat flux limit of 10 mW/cm^2^, assuming heat flux from both faces, minus the FPGA static power. For each BP and FPGA dimension pair, the ratio of how many more channels can be fitted on-implant with compression is also given.

### 4.2 Effects of Different Firing Rates and Communication Energy

It is worth noting that the BRs are taken from 3 datasets that all target the motor cortex during hand kinematic tasks, using Utah arrays. It may be that different tasks, cortical regions and microelectrode arrays will have varying FRs, which will affect the measured BRs. Similarly, a 20 nJ/bit communication energy estimate is used, but this could be overly optimistic. Such values have been observed outside the neural medium, and communication energies within the neural medium are likely to be much higher. As such, while the processing power and resource estimates in this work are robust, the communication energy, made up of the BR and comm. energy per bit, may vary.

Cortical areas, microelectrode arrays and tasks that have higher MUA FRs will be better served, all else held equal, by the windowed scheme. This is because the windowed scheme works better if the FRs are non-sparse. However, higher communication energies will make the communication power dominate, and so the importance of the processing power, which is higher in the DED encoding, will be less important when choosing a system. As such, it may be that, even with higher FRs than shown in this work, the DED scheme will be the best option given that it generally has lower BRs than the windowed scheme. This is especially true at finer BTRs.

Additionally, at higher communication energies, the role of data compression as a whole becomes more important. As such, the ratio of compressed to uncompressed channels that can be fit on-implant within heat limits, shown in Fig. 4, may be a conservative estimate. With significantly higher communication energies, a much larger ratio of compressed channels to uncompressed channels could be fit on-implant.

### 4.3 AH vs. SH encoding

While it is not shown in the main manuscript to simplify it, this work did not just look at SH encoding. For each of the four compression architectures, we also investigated modes with no lossless compression, with Adaptive Huffman (AH) encoding which trains the Huffman encoders using the data collected on-implant in real time, and the entropic bandwidth which gives the minimum BRs that could be achieved for each architecture given a perfect compression scheme. These modes are described in detail in the Supplemental Material, along with the BR results.

An interesting observation was that the SH encodings performed virtually identically to the AH encodings, which were fully adaptive and assumed perfect knowledge of the to-be-compressed data. As such, we can conclude that SH encoders are a remarkably hardware-efficient and well-performing compression scheme for MUA data. This is because the shape of the to-be-compressed data is more or less known *a priori*, and so on-implant training is unnecessary.

### 4.4 Sample Histograms for Mapping

The use of sample histograms to add adaptivity into the SH encodings, as in [14], was also investigated. Also to simplify the manuscript, detailed discussion and the BR results are given in the Supplemental Material. It was found that sample histograms did improve the BRs, but that their hardware and processing power costs, while not very large, were noticeable. It was deemed that, for a 20 nJ/bit communication energy, the use of sample histograms was not justified for the reduction in communication power. However, with larger communication energies, the use of sample histograms may be warranted to reduce the BR and so the communication power, which may be the dominating factor.

## 5 Conclusion

This work looked at various algorithms for compressing MUA data for WI-BMIs. It made significant use of SH encoders because of their hardware efficiency and good compression performance. This work found that the SH encoders performed exceptionally well, and that the MUA signal could be compressed to varying degrees depending on BP. For example, at the standard BP of 1 ms, between 4.6 and 26 times more channels can be fitted onto the same mm-scale implant merely because of the addition of the windowed and DED compression schemes analysed in this work.

It was found that for BPs ≤ 10 ms, the DED method had the lowest total power and reduced the BR by up to almost an order of magnitude relative to the classical windowed method (e.g. to approx. 151 bps/channel for a BP of 1 ms and 1000 channels on-implant.). However, at larger BPs the windowed method performed best (e.g. approx. 29 bps/channel for a BP of 50 ms, independent of channel count).

As such, this work can guide the choice of MUA data compression scheme for BMI applications, where the BR can be significantly reduced in hardware efficient ways. This enables the next generation of wireless iBMIs, with small implant sizes, high channel counts, low-power, small hardware footprint, and good temporal resolutions. MUA is already the most popular neural signal for WI-BMIs [3–5, 32]. The results from this work suggest that the MUA signal can be made even more attractive, given the possibility for hardware-efficient, ultra-low power compression. All code and results in this chapter have been made publicly available at [34].

