## Supplemental Material for "Ultra Low Power, Event-Driven Data Compression of Multi-Unit Activity"

November 25, 2022

#### 1 Calculating Bit Rates for Each Encoding

Here we give detailed explanations of each encoding, as well as how their Bit Rates (BR) are calculated. The BR notation is $\mathrm{BR}_{\mathrm{X-y}}$, where X is the encoding version, e.g. Basic (B), Static Huffman (SH), Adaptive Huffman (AH) or Entropic Bandwidth (E), and where y is the encoding, e.g. Windowed (W), Explicit Event-Driven (EED), Delta Event-Driven (DED), or Group Event-Driven (GED).

##### 1.1 Windowed encoding

###### 1.1.1 Basic implementation

The basic implementation of each encoding is implemented without any lossless compression. As such, the BR for the basic implementation of the Windowed encoding is given in Eq. 3 of the main manuscript.

###### 1.1.2 Static Huffman

Firstly, the average probability distribution of MUA FRs across channels, $\bar{p_i}$, was calculated. It is given by:

$$\bar{p_i} = \frac{1}{n} \sum_{j=1}^{n} p_{j,i} \tag{1}$$

where $p_{i,j}$ is the measured probability of a FR of $i$ MUA events/bin occurring on channel $j$ in a given BP across all of the considered data. As each channel's histogram had the same number of samples, $\bar{p_i}$ is proportional to the sum of the channels' histograms, meaning it is representative of the multi-channel data. $p_{i,j}$ was calculated for each considered number of channels $n$, BP and $S$, using a random selection of channels.

As such, the average BR per channel is given by:

$$\mathrm{BR}_{\mathrm{SH-W}} = \frac{\sum_{i=0}^{S-1} (CLVw - she_i \times \bar{p_i})}{n \times BP} \quad \text{[bps/channel]} \tag{2}$$

where $CLVw - she_i$ is the codeword length for FR = $i$ events/bin, where the codeword is given by the static Huffman encoder trained on a decaying exponential probability vector of length $S$, where FRs of 0 to $S-1$ events/bin are encoded.

###### 1.1.3 Adaptive Huffman

For the Adaptive Huffman (AH) version, a single Huffman encoder was trained on $\bar{p_i}$. Using $\bar{p_i}$ meant that each channel shared the same encoder. As such, it differs from the previous implementation in that the encoder is trained on the to-be-compressed data, whereas the previous encoder was trained on a decaying exponential, assumed to be a good fit to the data. The resulting BR is given by:

$$\mathrm{BR}_{\mathrm{AH-W}} = \frac{\sum_{i=0}^{S-1} (CLVw_i \times \bar{p_i})}{n \times BP} \quad \text{[bps/channel]} \tag{3}$$

where $CLVw_i$ is the $i^{th}$ element of the codeword length vector for the windowed encoding, which is a vector of the Huffman code lengths where the $i^{th}$ element corresponds to the value with probability $\bar{p_i}$.

##### 1.1.4 Entropic Bandwidth

The minimum number of bits per channel per second required for the windowed encoding is given by the average of the channels' Shannon entropies [1] divided by the BP:

$$\mathrm{BR}_{\mathrm{E-W}} = -\frac{1}{n \times \mathrm{BP}} \sum_{j=1}^{n} \sum_{i=0}^{S-1} p_{j,i} \times log_2\left(p_{j,i}\right) \qquad \text{[bps/channel]} \quad (4)$$

Here we assume each channel $j$ has its own encoder with codeword lengths $-log_2\left(p_{j,i}\right)$, giving the best compression possible for this encoding.

#### 1.2 Explicit event-driven encoding

##### 1.2.1 Basic implementation

In the windowed architecture, the FR per channel is encoded. The channel ID is implicitly encoded in bit position. To the best of the authors' knowledge, the following event-driven architectures are proposed for the first time in this work. In these event-driven architectures, the channel ID is explicitly encoded. E.g., in this 'explicit event-driven' architecture, the channel ID is only sent out if a non-zero FR of $i$ events/bin occurs on that channel, followed by a binary codeword representing $i$. If a channel has a FR of 0, that channel is missing entirely from the communicated codeblock. As such, although the channel ID has to be explicitly encoded, this is only if the FR is not 0 for that channel in the bin. For sparse signals, i.e. where FRs not equal to 0 are rare, this may offer improved compression over the windowed paradigm, where even FRs of 0 need to be communicated for each channel. As with the windowed architecture, the number of encoded FRs can be saturated at an integer value $S$.

As the channel IDs need to be explicitly encoded, the simplest method is to give the channels a standard binary codeword of length $k_1$, where:

$$k_1 = ceil(log_2(n)) \qquad \text{[bits]} \quad (5)$$

Similarly, the MUA FR for each channel is encoded as a binary codeword of length $m_2$:

$$m_2 = ceil(log_2(S-1)) \qquad \text{[bits]} \quad (6)$$

This differs from $m'$ in that a codeword of length $m'$ can only encode up to $2^{m'} - 1$ FRs, whereas a codeword of length $m_2$ can encode up to $2^{m_2}$ FRs. This is because the case of FR = 0 does not need to be encoded for the explicit event-driven encoding. An exception is if $S = 2$, i.e. $m_2 = 1$ and $i$ is limited to 0 and 1. In which case the $m_2$-length vector is unnecessary as the occurrence of the non-zero FR = 1 is already encoded in the sending out of the channel ID.

An example of the basic implementation, with $n = 4$, $k_1 = 2$ and $m_2 = 2$, is given by:

0001 0100 1011

This shows that channel 1 (00) had a FR of 2 events (01) in the bin, channel 2 (01) had FR = 1 (00), channel 3 (10) had FR = 4 (11), and channel 4 had a FR = 0 (absent). In this work, this encoding was named the explicit event-driven encoding because the FR per channel is explicitly encoded in the $m_2$-length codeword that follows the channel ID.

For calculating the BR, one only sends out the channel ID and FR codeword if a non-zero FR occurs on that channel in the given bin. As such, one sends out the channel ID of channel $j$ and its $m_2$-length codeword with probability $pc_j$:

$$pc_j = 1 - p_{j,0} \quad (7)$$

where $p_{j,0}$ is the probability of FR = 0 on channel $j$ in the average time bin at the given BP. As such, the BR is given by:

$$\mathrm{BR}_{\mathrm{B-EED}} = \frac{1}{n \times \mathrm{BP}} \sum_{j=1}^{n} pc_j\left(k_1 + m_2\right) \qquad \text{[bps/channel]} \quad (8)$$

##### 1.2.2 Static Huffman implementation

In this implementation for the explicit event-driven method, we used a SH encoder to losslessly compress the FRs. Only the FRs are compressed, whereas the channel IDs are not. This is very simple to implement in hardware while likely giving good compression. It uses the same $k_1$ length codeword for the channel IDs, but uses varying length codewords for the MUA FRs. As in the SH windowed implementation, the FR encoder was trained on a decaying exponential.

The BR is given by:

$$\text{BR}_{\text{SH-EED}} = \frac{1}{n \times \text{BP}} \sum_{j=1}^{n} \left( pc_j \times k_1 + \sum_{i=1}^{S-1} (CLVshe_i \times p_{j,i}) \right) \qquad \text{[bps/channel]} \quad (9)$$

where $CLVshe_i$ is the codeword length of FR = $i$, given by the static Huffman encoder trained on a decaying exponential probability vector of length $S-1$, where FRs of 1 to $S-1$ events/bin are encoded. Importantly, there is no FR codeword for $i = 0$ in this encoding, and the shortest codeword length is $CLVshe_1$.

##### 1.2.3 Adaptive Huffman implementation

In this AH implementation, we seek to compress both the FRs and the channel IDs, using the correct probability distributions observed on-implant. As such, this architecture differs from the previous one in that both the FR and the channel IDs are losslessly compressed. The prefix code nature of Huffman encoding can be taken advantage of to have two separate encoders: the first encoding the channel IDs, the second encoding the FRs. This is possible since we know the encoded FR will always follow the encoded channel ID, and so we know which encoder to use at each codeword. A design choice was made to train the FR encoder based on the average channel histogram, rather than have a separate encoder per channel. This is much more hardware efficient.

The ideal codeword length assigned to a FR depends on the relative probability of sending out FRs. As such, the FR encoder is trained on the relative probabilities $\widehat{pe_i}$:

$$\widehat{pe_i} = \frac{\sum_{j=1}^{n} p_{i,j}}{\sum_{i=1}^{S-1} \left( \sum_{j=1}^{n} p_{i,j} \right)}, \; i > 0 \quad (10)$$

where $\widehat{pe_i}$ represents the relative probability of any $i > 0$ FR occurring relative to other $i > 0$ FRs, averaged across all channels. It is similar to $\bar{p_i}$, the difference being that the case of $i = 0$ is discounted.

The use of relative probabilities to train encoders warrants spending some time on, because the motivation for absolute vs. relative probabilities is something that sets the considered event-driven architectures apart from the windowed one.

Relative probability gives how likely each symbol is to show up relative to other symbols, which is what encoders are trained on so as to give the ideal codeword length to each symbol. However, this becomes strange when it is no longer guaranteed that each channel will output a symbol that will receive a codeword. In this case, FRs = 0 are not encoded. To give the ideal codeword length, we should not encode the case where the codeword length will always be zero. Additionally, the sum of probabilities used to train the Huffman encoder should sum to 1. As such, we train the encoder on the relative probability of each non-zero FR occurring, so as to not waste bits on codewords that will never be used, e.g. FR = 0.

However, ultimately, the probability of that FR occurring is unchanged. Regardless of its optimal codeword length, it will occur at the frequency it occurs at. As such, we need to differentiate between the absolute probability, which predicts how often the FR will occur, and the relative probability, which gives the ideal codeword length for the FR. And so, for the average FRs across channels, we differentiate between $\bar{p_i}$ and $\widehat{pe_i}$.

The same concept applies to the channel IDs. Some channels may be dysfunctional and always have FRs of 0, in which case they should not be encoded. While unlikely, it is ideal in terms of compression to account for it. The relative probability $\widehat{pc_j}$ of sending out channel $j$'s ID is given by:

$$\widehat{pc_j} = \frac{pc_j}{\sum_{l=1}^{n} (pc_l)} \quad (11)$$

The BR is then given by:

$$\text{BR}_{\text{AH-EED}} = \frac{\frac{1}{n} \sum_{j=1}^{n} (CLVc_j \times pc_j) + \sum_{i=1}^{S-1} (CLVfr_i \times \bar{p_i})}{BP} \qquad \text{[bps/channel]} \quad (12)$$

where $CLVc_j$ is the Huffman codeword length of the $j^{th}$ channel ID, and $CLVfr_i$ is the Huffman codeword length for the $i$ events/bin MUA FR, trained on $\widehat{pe_i}$. The right hand side concerning $CLVfr$ does not need to be divided by $n$ because the channels have already been averaged in $\bar{p}_i$.

###### 1.2.4 Entropic Bandwidth

To calculate the lower limit of how many bits are needed for the explicit event-driven encoding, four aspects are calculated. The first two are the absolute and relative probability of channels having FR $> 0$, i.e. $pc_j$ and $\widehat{pc_j}$ as calculated in Eq. 7 and 11.

The second two are the absolute and relative probabilities of each FR per channel, $p_{i,j}$ and $\widehat{p_{i,j}}$. $\widehat{p_{i,j}}$ is different from $p_{i,j}$ in that the case of a $i = 0$ FR is not represented. As such, $\widehat{p_{i,j}}$ is given by:

$$\widehat{p_{i,j}} = \frac{p_{i,j}}{\sum_{i=1}^{S-1} p_{i,j}}, i > 0 \tag{13}$$

In the entropy calculation, each channel was considered to have its own unique encoder, which is ideal in terms of compression if not hardware realisation. This is why we used $\widehat{p_{i,j}}$ instead of $\widehat{pe_i}$, the latter of which is averaged across channels.

From these four elements, we calculated the entropic BR, which is the minimum average number of bits per channel per second required to represent the channel IDs and the subsequent $i > 0$ FRs.

$$\text{BR}_{\text{E-EED}} = -\frac{1}{n \times \text{BP}} \sum_{j=1}^{n} \left( pc_j \times log_2(\widehat{pc_j}) + \sum_{i=1}^{S-1} (p_{i,j} \times log_2(\widehat{p_{i,j}})) \right) \text{ [bps/channel]} \tag{14}$$

#### 1.3 Delta-event-driven encoding

##### 1.3.1 Basic implementation

It may be that, rather than communicating out discrete codewords for the channel IDs, it would be better to send out the encoded difference between adjacent channel IDs. In effect, we are delta-sampling the communicated out channel IDs, i.e using a run-length encoding. If a channel has a FR above 0, it is given a $\Delta$ value by subtracting the ID of the previous channel to have a FR above 0, $j_{previous}$, from the current channel's ID, $j_{current}$. I.e., $\Delta = j_{current} - j_{previous}$. The first channel ID to be sent out in the BP will be $\Delta$-sampled relative to a channel ID of 0.

However, there is no clear basic implementation of this method that does not involve compressing the delta-samples. This is because the amount of bits required to represent each delta-sampled channel ID $\Delta \in [\mathbb{Z}, 1 \leq \Delta \leq n]$ is the same $k_1$ bits as required to represent the channel IDs directly. This is because the most extreme case of $\Delta = n$, where only the last channel has a FR above 0, needs to be accounted for. Therefore $ceil(log_2(n)) = ceil(log_2(n)) = k_1$ bits are required to encode $\Delta$. As such, the basic implementation of this method was ignored and we moved straight the SH implementation.

##### 1.3.2 Static Huffman implementation

In this implementation, we assigned a SH codeword to the $\Delta$-sampled channel IDs, where smaller $\Delta$ values received shorter codewords. This way, if channel IDs are communicated out often, the $\Delta$ codewords will likely be shorter than the standard $k_1$ bits.

###### 1.3.2.1 Training the $\Delta$ encoder

However, a design choice has to be made on how to train the delta-sampled channel ID encoder. It was decided to train the decoder on a decaying exponential so that smaller $\Delta$ values were given shorter codewords. However, the ideal rate of decay is an unknown parameter. Larger rates of decay will give smaller differences shorter codewords, which is ideal if channel IDs are communicated out often. Smaller rates of decay will give more equal codeword lengths for different $\Delta$ values. This is ideal if the$\Delta$ values vary significantly. As such, a set of $d = -10^f$ rates were considered, where $d$ is the exponent of a decaying exponential and $f$ is an integer with $f \in [\mathbb{Z}, -3 \leq f \leq 2]$. As such, decaying exponents of $-10^{-2}$ to $-10^{3}$ were considered. The SH encoder for delta-sampled channel IDs was then trained on a probability vector $p_d(d, \Delta)$ of length $n$, where:

$$p_d(d, \Delta) = \frac{e^{d \times \Delta}}{\sum_{g=1}^{n-1} e^{d \times g}} \tag{15}$$

As shown in Eq. 15, $p_d(d, \Delta)$ was normalised across all $\Delta$ so as to sum to 1 for each $d$ value. For each BP, $S$ and $n$ combination, each considered decay exponent $d$ was used to train a delta-sampled channel ID static encoder. The channel IDs were then delta-sampled. $\Delta$ was then encoded using the static encoder trained on $p_d(d, \Delta)$. The BR was stored for each $d$, and the best performing $d$ was then chosen for each BP, $S$ and $n$ combination.

###### 1.3.2.2 Saturating at a $\Delta_{max}$ value

However, having an on-implant Huffman encoder stored in a Look-up Table (LUT), with a unique codeword for each $\Delta$ value for large $n$, can have significant memory requirements. As such, it was analysed whether a shorter codebook of $\Delta$ values could be used, truncated at a $\Delta_{max}$ value. If a $\Delta > \Delta_{max}$, then a reset signal, equal to $\Delta_{max} + 1$, was communicated out along with how many times the reset signal should be read and the remainder, $\Delta_{max} - \Delta$. As such, $\Delta$ was transformed into a concatenated sequence of:

$$\Delta \rightarrow \Delta_{max} + 1,\ \gamma,\ B$$

where $\Delta_{max} + 1$ is the reset signal, $\gamma$ is an integer value expressing how many times we should read the reset signal, and $B$ is the remainder. For example, for $\Delta_{max} = 10$ and $\Delta = 45$:

$$\Delta \rightarrow 11, 4, 5$$

where we first communicate the reset signal indicating that $\Delta_{max}$ has been exceeded ($\Delta_{max} + 1 = 11$), then we communicate how many times it has been exceeded ($\gamma = 4$), and finally the remainder ($B = 5$).

The $\Delta_{max} + 1$ reset value was encoded using the delta-encoder trained on $p_d(d, \Delta)$, and the quotient value $\gamma$ and remainder $B$ were given fixed-length codewords of lengths $\theta_\gamma$ and $\theta_B$ respectively. Since the fixed-length codewords only ever occur after the reset signal, the sequence is fully decodable.

This saved memory, since there were fewer possible $\Delta$ codewords. However, limiting $\Delta$ to $\Delta_{max}$ in this way increased the processing hardware resources and power. $\Delta_{max}$ values of $\Delta \in [2^{[3,\,6,\,7,\,8,\,9,\,10]},\ n]$ were considered. If for a given parameter combination, $n < \Delta_{max}$, that parameter combination was ignored. The values $\gamma$ and $B$ are given in the Table 1.

##### 1.3.3 Adaptive Huffman implementation

For the AH version, no $\Delta_{max}$ was considered. Similar to the SH version, for the AH version the data was iterated through time-wise and the $\Delta$ values stored across all timesteps and channels. In effect, we counted the number of occurrences of each $\Delta$ value, and stored them in a vector $u\Delta$. A probability vector $\widehat{p_\Delta}$, of length $n$, was then obtained that gave the relative probabilities of each $\Delta$ value occurring throughout the data:

$$\widehat{p_\Delta} = \frac{u\Delta}{\sum_{\Delta=1}^{n} u\Delta} \tag{16}$$

$\widehat{p_\Delta}$ was used to train a Huffman encoder, that gave the encoded codeword lengths $CLV_\Delta$ for each $\Delta$ value. The BR is given by:

$$\text{BR}_{\text{AH-DED}} = \frac{\sum_{\Delta=1}^{n}(CLV_\Delta \times u\Delta)}{n \times v/\text{BP}} + \frac{\sum_{i=1}^{S-1}(CLVfr_i \times \bar{p_i})}{\text{BP}} \quad \text{[bps/channel]} \tag{17}$$

where $CLV\Delta_i$ is the codeword length of the $\Delta$ values given by the Huffman encoder trained on $\widehat{p\Delta}$. $v$ is the data length in seconds, i.e. $v$ = 100 s.

On the left hand side of Eq. 17, we can see that we take the codeword length weighted sum of all of the occurring delta values, and divide by the total data length in samples, i.e. $n \times 100\,\text{s/BP}$, to get the average bps/channel for the channel IDs. We combine this with the FR component on the right hand side, which is the same as in Eq. 12, where we train a single AH encoder for all of the channels.

Table 1: Values for $\gamma$, $B$ and $\theta$ values for $\Delta_{max}$ in the SH Delta Event-Driven encoding. 'floor' is the floor operation, and 'mod' is the modulus.

| Parameter | Value |
|---|---|
| $\gamma$ | floor(($\Delta$ - 1) / $\Delta_{max}$) |
| $B$ | mod($\Delta$ - 1, $\Delta_{max}$) + 1 |
| $\theta_\gamma$ [bits] | ceil(log$_2$(($n$ / $\Delta_{max}$) - 1)) |
| $\theta_B$ [bits] | ceil(log$_2$($\Delta_{max}$)) |

##### 1.3.4 Entropic Bandwidth

There is a minor point to mention in the entropic calculation for the explicit event-driven method with delta-sampled channel IDs. We assume that, unlike in the AH version above, each channel has its own unique FR decoder. This is optimal for compression but not for hardware complexity. This is theoretically possible as the channel ID is encoded as delta-samples, and from the delta-sample the actual channel ID is derived off-implant. Having derived the channel ID off-implant, one can decode the channel's FR using the channel's FR decoder.

The BR is given by:

$$\mathrm{BR}_{\mathrm{E-DED}} = -\frac{\sum_{\Delta=1}^{n}(u\Delta \times log_2(\widehat{p\Delta}))}{n \times v/\mathrm{BP}} - \frac{\sum_{j=1}^{n}\sum_{i=1}^{S-1}(p_{i,j} \times log_2(\widehat{p_{i,j}}))}{\mathrm{BP}} \quad \text{[bps/chan]} \quad (18)$$

#### 1.4 Group event-driven encoding

##### 1.4.1 Basic implementation

In this group event-driven encoding, one uses position and stop symbols to encode the FRs. As in the explicit event-driven encoding, one explicitly encodes the channel ID, but here one encodes the FR per channel implicitly in channel ID position. For example, in decimal,

$$2\,4\,stop\,1\,6\,stop\,stop\,3$$

signifies that channels 2 and 4 had a FR of $i = 1$ in the given bin, channels 1 and 6 had a FR of $i = 2$, channel 3 had FR $i = 4$, and the rest of the channels had FR $i = 0$. As with the explicit event-driven encoding, this event-driven encoding benefits from not having to encode channels with FRs of 0 in the given bin, making it perform well for sparse signals. As with the other architectures, the measurable FRs can be saturated at $S$.

There is a design choice to be made on whether each number $i \geq 2$ of events should have its own stop codeword, or whether all event numbers should share the same stop codeword. We opted to have all of the transitions have their own codeword. This alleviates the risk of requiring multiple stop codewords next to each other. This risk is shown in the example above where no channels had 3 events/bin but one had 4. As such, all values to the right of the $i$ stop symbol have FRs $\geq i$, where $i \geq 2$. E.g., channel IDs before the $i = 2$ stop symbol have FRs equal to 1, and there is no $i = 1$ stop symbol.

The stop symbol for each $i \geq 2$ has, in the average codeblock, a probability $ps_i$ of occurring. $ps_i$ is equal to the probability that there will be at least 1 channel on which there is a FR of $i$. This is equal to the complement of the probability of no channels having a FR of $i$. As such, $ps_i$, a vector of length $S - 2$, is given by:

$$ps_i = 1 - \prod_{j=1}^{n}(1 - p_{j,i}), \quad i \geq 2 \quad (19)$$

For the codeword lengths, the simplest implementation is to give the stop symbols and the channel IDs a length of $k_2$ bits, where:

$$k_2 = ceil(log_2(n + S - 2)) \quad \text{[bits]} \quad (20)$$

E.g. $n = 2$, $S = 4$, means that there are 2 channels, FRs between 0 and 3 inclusive can be encoded, and $k_2 = 2$. Channel 1 gets a codeword of 00, channel 2 gets a codeword 01, and the stop symbols for $i = 2$ and $i = 3$ get codewords of 10 and 11 respectively. It warrants mentioning that for large numbers of channels, typical $S \leq 20$ values will have relatively little impact [2]. As such, relatively large dynamic ranges may be achievable will little to no cost in this basic group event-driven architecture for large $n$. Alternatively, the number of channels should be chosen carefully in conjunction with $S$ so as to maximize the use of the range given by $k_2$.

The average BR per channel is given by:

$$\mathrm{BR}_{\mathrm{B-GED}} = \frac{1}{n \times \mathrm{BP}}\left(\sum_{j=1}^{n} pc_j \times k_2 + \sum_{i=2}^{S-1} ps_i \times k_2\right) \quad \text{[bps/channel]} \quad (21)$$

##### 1.4.2 Static Huffman implementation

The group event-driven encoding does not have an obvious SH version. This is because the codeword lengths are all relative to both channel ID and stop symbol frequency. While these could be estimated, e.g. the channels all have an equal likelihood of occurring and the stop symbols follow a decaying exponential, it becomes a lot of guesswork very quickly in terms of the relative probabilities. As such, no SH version was considered for the group event-driven encoding. The basic version, where all codewords had an equal length, was assumed to be close to what a SH version would have achieved.

##### 1.4.3 Adaptive Huffman implementation

In the explicit event-driven architecture, one knows that the channel ID codeword is followed by a codeword encoding the channel's FR. However, in this architecture it is unknown whether the next codeword will be a channel ID or a stop symbol. As such, the codewords need to uniquely decodable, and so must come from the same encoder. This is also why the channel IDs and stop symbols shared the same binary $k_2$ codeword basis in the basic implementation. The Huffman encoder in the group event-driven architecture must be trained on the combined probabilities of the channel ID and stop symbol occurring.

$$pg = [pc,\, ps] \tag{22}$$

where $pg$ is the collated probability vector of $pc$ and $ps$. $ps_0$ and $ps_1$ are not included in the collation as they are not valid values (see Equation 19). The relative probabilities are given by:

$$\widehat{pg_a} = \frac{pg_a}{\sum_{a=1}^{n+S-2} pg_a} \tag{23}$$

where $a \in [\mathbb{Z},\, 1 \leq a \leq n+S-2]$ is the index of the concatenated probability vector. The encoder was trained on $\widehat{pg}$. The BR is given by:

$$\text{BR}_{\text{AH-GED}} = \frac{\sum_{a=1}^{n+S-2}(CLVg_a \times pg_a)}{n \times \text{BP}} \quad \text{[bps/channel]} \tag{24}$$

where $CLVg_a$ is the length of the Huffman codeword associated with the $a^{th}$ element of the combined probability vector $pg$. Our convention was that if $a \leq n$, $CLVg_a$ represents the length of a channel ID codeword. Otherwise, it represents the length of a stop symbol.

##### 1.4.4 Entropic Bandwidth

The average entropic BR for each channel for the group event-driven encoding is given by:

$$\text{BR}_{\text{E-GED}} = -\frac{1}{n \times \text{BP}} \sum_{a=1}^{n+S-2} (pg_a \times log_2(\widehat{pg_a})) \quad \text{[bps/channel]} \tag{25}$$

#### 1.5 Group event-driven encoding with delta-sampled channel IDs

A version of the group event-driven encoding with delta-sampled channel IDs was not considered. This is because one would have to delta-sample the channel IDs, but would have to delta-sample the channel IDs within the same FR for the delta values to have any sense. E.g.

$$3\;1\;stop_2\;5\;1$$

would indicate that channels 3 and 4 had FR of 1, and that channels 5 and 6 had FRs of 2. Adding in variable length codewords for the combined relative probabilities of each delta-value and stop symbol seemed to be very overly complicated an encoding, frankly. It is theoretically possible, but a static pre-trained version seemed to be a significant amount of guesswork relative to the relative probabilities, and the hardware implementation was not attractive to the authors. As such, although perhaps feasible as an encoding, a group event-driven encoding with delta-sampled channel IDs was not considered further by the authors.

#### 1.6 Sample Histogram for Firing Rate mapping

The MUA histogram, for BP $\leq$ 100 ms, typically follows a decaying exponential. In other words, smaller FRs are more common than larger ones. However, this is not always the case. As such, automatically assigning shorter codewords to smaller FRs will not always give optimal compression. In [2], the use of a sample histogram was examined to address this problem. The beginning of each recording was used to fill a sample histogram. This histogram was then used to estimate the relative frequencies of the FRs for each channel. The most common FRs in the histogram were then, for the rest of the data in each channel, assigned the shortest codewords via sorting. This was referred to as mapping the most common FRs to the shortest codewords, given the sample histogram estimate. As such, some semi-adaptability was introduced into the SH encoders. This process is shown in Fig. 1.

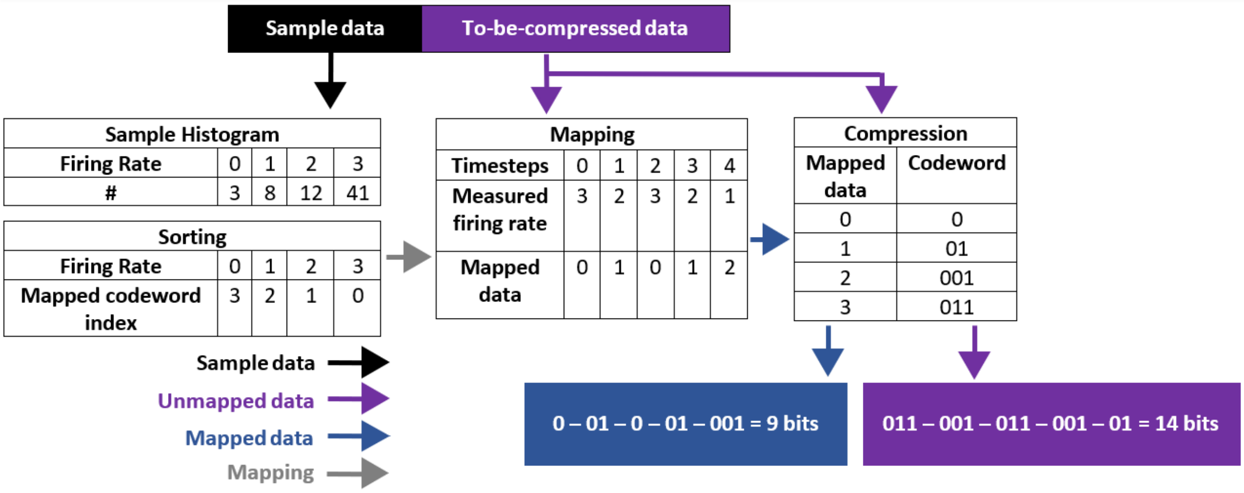


Figure 1: Use of a sample histogram to improve bit rates. A sample histogram is derived from the beginning of each channel's recording. It is then sorted using a hardware-efficient sorting, where smaller indices are given to more common firing rates. The sorting is stored as a mapping, used to sort the rest of the data, i.e. the to-be-compressed data. The data is then compressed after mapping, where if a firing rate was found to be the $x^{th}$ most common in the sample histogram, it was given the $x^{th}$ shortest codeword. As such, the data histogram is approximated by taking a sample, and if the sample is well-representative of the rest of the data, this may help shorter codewords be given to more common firing rates, improving compression. In the example we can see the mapped compressed data requires only 9 bits, relative to the unmapped compressed data which requires 14 bits.

Using a histogram for mapping the firing rate was extremely hardware efficient (for small $S$), since the sorting procedure was implemented using only combinatorial logic [2]. Furthermore, the sample histogram and sorting logic modules were shared across channels, with access to them multiplexed.

As such, for the SH and AH implementations of each encoding, we looked at the effect of including a sample histogram. We did not use them for the basic implementations, because there was no lossless compression and all codewords are in equal length. We considered histogram bin sizes of $hs$ = 0, 2, 3, 4, 5, and 6 bits. In the case of 0 bits, no histogram, sorting or mapping was used. Otherwise, the beginning of each channel's recording was used to train the histogram. When $2^{hs}$ FR samples had been measured by the histogram for a channel $j$, the histogram training was ended for channel $j$. It was then sorted, and the mapping produced. The rest of the data was then compressed, after mapping. As such, only $v/\text{BP} - 2^{hs}$ samples were compressed in the histogram-included versions of these encodings, where $v = 100\,\text{s}$ is the total length of each channel's recording.

For the event-driven encodings, the effect is very important. With the FR histogram and mapping, one is no longer sending out the information from a channel $j$ if it has a FR $> 0$. Rather, one is sending out its information if it has a FR different from its most common FR, as estimated by the histogram. This can greatly improve the compression if a channel's most common FR is not 0.

#### 2 Hardware design

The AH implementations require us to generate the Huffman codebook on implants, and assume perfect knowledge of the data to be compressed, which is not achievable in practice. Therefore, no hardware implementation was done for the AH encodings.

The windowed, explicit event-driven and delta-event-driven implementations share a similar hardware architecture. The SH delta-event-driven encoding is considered as the 'full version' architecture, while the explicit event-driven and windowed architectures are pruned versions. As such, for conciseness, we introduce the implementation of the delta-event-driven encoding and describe how the other architectures can be derived from these.

#### 2.1 Delta-event-driven encoding

This compression architecture consists of 7 components. Four of them have been described in detail in previous work [2]. These four are the binner, histogram, mapper and encoder. As such, we only give a brief description of them.

- The binner is a counter that counts the number of detected spikes (FR) at the given BP.
- The histogram accumulates the frequency of different FRs to identify the most common FR for each channel. This is required for sorting and mapping.
- The mapper is the module responsible for assigning each FR to its codeword. The mapper maps the most frequent FR, as determined in the histogram, to the shortest codeword and maps the other FRs accordingly with a pre-defined combinatorial logic that defined the sorting. As detailed in Section 2.6 of the main manuscript, this mapping aligns the real-time FR distribution with the distribution that the Huffman encoder is trained on, which helps maximise the on-implant compression performance.
- The encoder is a LUT that encodes the mapped FR into a Huffman codeword.

In order to achieve the event-driven encoding and delta-event-driven encoding, two more components have been designed. The first component is the comparator. This compares the FR to be compressed with the most frequent FR as measured by the histogram. If no histogram/sorter/mapper is used, then the most common FR is taken to be 0. The comparator ensures that only the not-most-common FRs are sent out. The second component is responsible for the channel delta-sampling. It calculates the difference of the current channel to-be-transmitted to the last transmitted channel. ROM is used to store the original channel ID codeword instead of using a LUT implementation. This saves on logic cell resources.

For implementing multi-channel compression with minimum resource usage, we follow the time-sharing architecture as in cite [2, 3]. All channels share the same processing components, and the signals of each channel are processed interchangeably. RAM is used to store the variables for each channel while the processing circuits are busy with another channel.

#### 2.2 Windowed encoding

The basic implementation of windowed encoding transmits the binned raw FR of all channels with no further operation. Its SH implementation compresses the binned FR with a pre-trained Huffman encoder. It also has the option to use a histogram and mapper to enhance the compression performance.

#### 2.3 Explicit Event-Driven encoding

The basic implementation of the explicit event-driven encoding uses the binner and comparator for obtaining non-zero FR and channel number for transmission. The SH implementation, similar to the windowed one, makes use of Huffman encoder to compress the FR. As with the windowed encoding, a histogram and mapper can be used to better than chances that the most common FRs are given the shortest codewords.

#### 2.4 Group Event-Driven encoding

The group event-driven encoding is different from the other three. The high-level summary is that the FR from each channel is read. The channel IDs with the same FR are then grouped. Once every channel has been read, the groupings/concatenations of channel IDs are sorted in order of FR with interleaved stop symbols to create the final bit stream.

In hardware, this consists of a sorter and a package generator. The sorter counts the FR in each channel continuously. Meanwhile, the multi-channel FRs are sorted in descending order. The package generator generates the bit stream based on the sorted FRs according to the policy described as section 1.4.

Sorting can be resource and time consuming, especially in hardware. Here it is implemented using a Finite State Machine (FSM). We sort the bin counts (FRs) in real time as they accumulate. Each time a spike is detected, only one channel's FR changes and it only increases by one. Swapping this FR with the furthest value in front which is smaller than it keeps the sorted FRs in order. To ensure that the FR and channel number are trackable after swapping, two arrays of registers are needed. The first stores the channel numbers of the sorted entry. The second stores the index number of each channel after swapping. Therefore, the temporal and spatial cost increases linearly with the channel

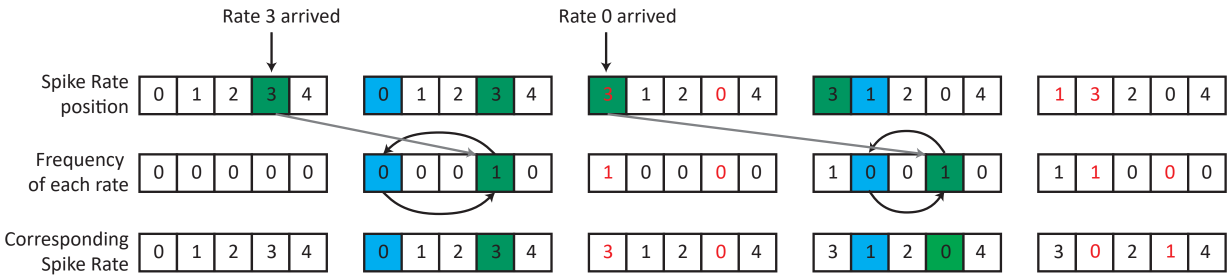


Figure 2: An example of the sorting algorithm. Fequency of each channel: sorted spike rate of each channel. Corresponding channel number: the channel number corresponding to frequency above. Channel position: the index of the channel corresponding to spike rate of each channel array. The workflow goes as below: A spike is first detected in channel 3, Channel 3 spike rate then increases by one. The number to be swapped for sorting is the furthest smaller rate left to the rate just increased (the green 1 in B), which is the 0 in blue. Swapping then happens in all three arrays. Another spike is detection in Channel 0. According to array A, the spike rate of channel 0 is stored in index 3 of array B. The number to be swapped is the most left 0 in B and all corresponding values in A, B and C are swapped then.

number, i.e. O($n$), where $n$ is the number of channels. This low complexity sorting is achieved by taking advantage of the nature of the binning where only one entry is increased by one at each spike detection. An example is give in fig.2, illustrating the sorting algorithm.

The package generator then scans the sorted FR while neglecting any zeros in the tails. The channel numbers are packaged/concatenated and the stop symbols are inserted whenever a new FR is scanned in the sorted FR sequence.

The group event-driven encoding however has more hardware complexity than the other three encodings. This is because the channel IDs must be placed in their correct location in the bit stream. This is the case even for the basic implementation. This is a significant hardware cost. Additionally, this sorted collation occurs at every BP and so adds a processing power burden, especially at larger channel counts as the channel ID codewords are longer. As such, although the group event-driven encoding generally outperformed the explicit event-driven encoding in terms of compression, its added processing power and hardware complexity may make it less practical as an encoding. It is however interesting that the entropic and basic implementations gave similar BRs, suggesting that the basic implementation gives almost ideal compression for the group event-driven encoding.

### 3 Hardware Results

There is basically no difference in hardware costs between the Basic and SH implementations of each encoding. As such, we will show only the SH hardware results.

#### 3.1 The number of logic cells is similar across encodings

Table 2 shows the hardware requirements for the SH Windowed (SH-W), Explicit Event-Driven (SH-EED) encodings. We also show the Delta Event-Driven (SH-DED) encodings as a function of channel count (averaged across all other parameters). This is because the SH windowed and SH explicit event-driven hardware requirements do not increase with channel count However, the number of logic cells for the SH-DED encoding does depend on channel count, thus why different channel count values are shown for the SH-DED encoding.

It can be observed that there is little difference in hardware requirements between the encodings even in the most extreme case of $n = 30000$ for the SH-DED encoding. Therefore, we can conclude that the choice of encoding is basically unaffected by the number of required logic gates, although we would have a slight preference towards the SH-W and SH-EED encodings.

#### 3.2 The memory requirements vary by encoding

Table 3 shows the trend of required resources for the SH-EED and SH-DED encodings as a function of $n$, $S$ and $\Delta_{max}$. Importantly, these trends are in addition to the required resources for the SH-W encoding. As such, the SH-EED and SH-DED encodings require more resources than the SH-W encoding, and the additional resources follow the trend shown in Table 3.

Table 2: Average Logic cell and power across different parameters of SH-W SH-EED, and SH-DED at different channel counts

| | SH-W | SH-EED | SH-DED n = 10 | SH-DED n = 100 | SH-DED n = 1000 | SH-DED n = 10000 | SH-DED n = 30000 |
|---|---|---|---|---|---|---|---|
| Logic cells | 230 | 237 | 258 | 271 | 290 | 303 | 308 |
| Power (uW) | 0.96 | 1.04 | 1.06 | 1.07 | 1.11 | 1.23 | 1.76 |

Table 3: Logic cell, RAM and ROM scale-up speed of SH-EED and SH-DED with different parameters. Importantly, these trends are relative to the SH-W encoding. As such, as they are positive values, they are in addition to that required for the SH-W encoding.

| | SH-EED | SH-DED |
|---|---|---|
| Logic cells | log(S) | log(S)+log(n) |
| RAM | nlog(S) | nlog(S) |
| ROM | nlog(n) | $\Delta_{max}$log($\Delta_{max}$) |

We can observe that the required logic cells increase as a function of channel count $n$ and $S$ in the SH-DED encoding, whereas for the SH-EED encoding it only increases as a function of $S$. However, the SH-DED encoding's memory requirements scale far better as a function of channel count $n$ than the SH-EED encoding, since one can set $\Delta_{max} << n$. As such, as far as memory is concerned, SH-DED scales better with channel count, assuming $\Delta_{max}$ is set to a value smaller than $n$.

As such, we can conclude that SH-W is the most hardware efficient option. The SH-EED and SH-DED encodings require additional logic cells and memory relative to the SH-W encoding. However, the requirement for additional logic cells is minor for the SH-EED and SH-DED encodings, especially considering these logic cells are shared via multiplexing across channels. Between the SH-EED and SH-DED encodings, SH-DED consumes significantly less memory if $\Delta_{max}$ is set to a value $\Delta_{max} << n$.

### 4 Hardware result analysis

#### 4.1 Hardware setting selection

Hardware cost includes the area occupation and power consumption. These two factors determine the suitability of one algorithm for on-implant use. However, the ultimate solution for on-implant signal processing should use ASIC design to minimise the cost, while we assessed the hardware cost using FPGA. Assessing using FPGA can reduce the development time significantly, meanwhile FPGA power consumption is proportional to the ASIC design and its resource usage is directly related to the area occupation. Considering the enormous parameter space involves in this work, we used FPGA resource usage and core dynamic power as a reference to assess the suitability of different proposed algorithms and compare across them.

To break down the hardware cost, the resource usage consists of the logic cells and block RAMs. Logic cells consists of LUTs and flip-flops constructing the combinational and sequential processing logics. The block RAMs can be used to construct RAM and ROM for storing temporal logic status or Huffman codewords. The power consumption consists of the static power and dynamic power, while different implementation only contributes to the dynamic power. For the ease of presenting, we further breakdown the dynamic power consumption into the power contributed from the circuit working at clock frequency and the circuit working at bin frequency. The bin frequency circuit share is much lower than that of the clock frequency circuit. The mapper and encoder work at bin frequency and only consumes 0.06 at $1ms$ bin period. Longer bin period leads to the power consumption less than 0.01 which is neglectable. Clock frequency power can be more significant and all power mentioned later will be the dynamic power per channel from the clock frequency circuit including binner, asyncing circuits, RAM and ROM.

Considering the superior performance of the SH over B implementation and the tiny cost of implementing the encoding (10% contribution to resources and neglectable effect on power consumption) comparing to AH, we prefer the SH architecture in this work. The group event-driven implementation consumes much more than others does (over 800 logic cells). Therefore, the best architecture should be selected among SH-W, SH-EED and SH-DED.

Both SH-EED and SH-DED are built upon the SH-W implementation. The average logic cell usage and power consumption is given in Table.2, SH-EED only takes less than 5% extra logic cells which is neglectable. SH-DED uses more logic cells especially when channel count increases, but it is still acceptable if we consider the average resource usage per channel. The power consumption shows similar tends as the logic cells (Note that the power rocketing from $n$ = 10000 to $n$ = 30000 is from the addition memory usage). Therefore, when SH-EED or SH-DED can provide better compression performance, they are preferred instead of SH-W because of the tiny cost.

In order to make selection between SH-EED and SH-DED, Table. 3 summarises how logic cells, RAM and ROM scales with different parameters. Details on how this table been derived is given in later two sections. One can notice that when the channel count increases, the logic cells of SH-DED increases in log-scale. However, the ROM usage of SH-EED increases in nlog-scale, which is much faster than the former one which grows with the $\Delta_{max}$. We therefore prefer SH-DED especially when channel count is extreme. Setting a small $\Delta_{max}$ value can effectively limits the aggressive growth of the circuit size.

Next comes to select a reasonable $\Delta_{max}$ value. However, there is no clear clue on how the number of extra logic cells and power consumption are varying with $\Delta_{max}$ the power consumption is overall consistent. Therefore, one should trade off their ROM/area availably and BR requirement when selecting $\Delta_{max}$. As the growing $\Delta_{max}$ has opposite effect on the ROM usage and BR reduction. Fig.3 shows the trend of power, bit rate and memory occupation (RAM+ROM) with different $\Delta_{max}$ values at bin period 1ms. 64, 128 and 256 are three sweet points to set $\Delta_{max}$. Around 30% and 45% bit rate reduction can be achieved respectively with acceptable memory/size cost. Power consumption in this case is not a concern as it stays at similar levels among settings. When bin period exceed 1ms, there is no bit rate gain when $\Delta_{max}$ is greater than 64. Therefore, SH-DED with $\Delta_{max} = 64$ is recommended when async-compression can outperform the sync-compression.

It was decided to fix $\Delta_{max}$ at a value for each BP and $n$ combination, which are the values that are relevant to $\Delta_{max}$. This was done by comparing the hardware results to the compression results for different $\Delta_{max}$. For each BP, the BR, total processing power and required memory for each $n$ and $\Delta_{max}$ were observed. A design choice was then made, to find a $\Delta_{max}$ value that minimises BR, processing power and memory. An example is shown in Fig. 3, for BP = 1 ms. As such, for a BP of 1 ms, we decided to set:

$$\Delta_{max} = \begin{cases} n, & \text{if } n \leq 100 \\ 64, & \text{if } n > 100 \end{cases} \tag{26}$$

For BPs larger or equal to 5 ms, it was found that there was little benefit to BR to increasing $\Delta_{max}$ beyond 64. As such, for BPs $\geq$ 5 ms, the following $\Delta_{max}$ values were set:

$$\Delta_{max} = \begin{cases} n, & \text{if } n \leq 10 \\ 64, & \text{if } n > 10 \end{cases} \tag{27}$$

This gave small values for $\Delta_{max}$, minimising the required memory as given in Table 3, while also giving good compression performance.

The resource usage of SH-W has been introduced in detail in [2], so here we here will only introduce the extra cost of SH-EED and SH-DED adding up to the SH-W later. As a reference, at bin period of 1ms, S = 3, histogram size = 2, SH-W occupies 129 logic cells and consumes 0.96$\mu W$. The idea is to investigate how the extra hardware cost scales with certain parameters, and whether the extra cost in hardware worth the gain in BR reduction in certain BP.

##### 4.1.1 SH-EED

The extra logic cell cost of SH-EED scales as $LC_{SH-EED} \propto log(S)$. In other words, in proportion to the bit width for $S$. However, comparing to the logic cell usage of the example SH-W implementation, this below 10 additional logic cell usage is neglectable.

When it comes to the memory usage, the RAM usage stays the same while it needs extra space of ROM to store the channel ID codewords. That ROM usage scales as $ROM_{SH-EED} \propto n \times log(n)$, as the bit width of the codeword scales logarithmically with the channel number.

As for the power consumption, our results suggest that the upscaling of the memory usage has less impact on power consumption, while the increasing logic cell usage does. Therefore, as the extra logic cells are neglectable, even though the memory occupation scales up quickly with the increasing channel count, the power consumption only increased for only 0.03 from 10 channels to 1000 channels.

##### 4.1.2 SH-DED

This architecture was built upon the SH-EED adding the logic counting the channel difference and the number of resetting. This extra logic is no longer neglectable. When there is no limitation on the max channel difference, i.e.

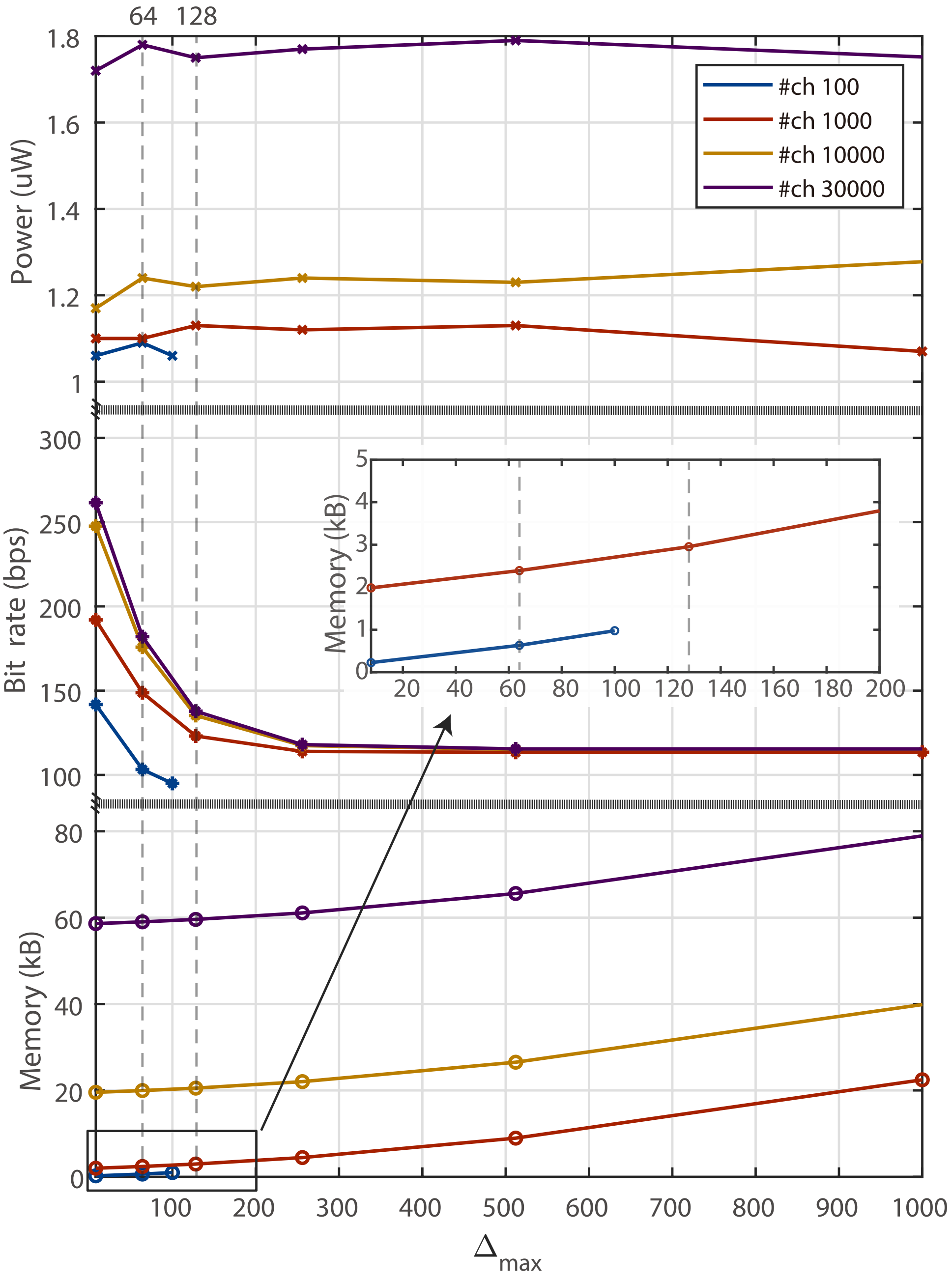


Figure 3: The processing power consumption, bit rate and memory usage of the FPGA implementation with different $\Delta_{max}$ values with BP = 1 ms. We can see that processing power consumption is largely unaffected by $\Delta_{max}$, and that channel count $n$ is the main consideration for processing power. $\Delta_{max}$ mainly affects BR and memory consumption, where memory increases roughly linearly with $\Delta_{max}$ and BR decreases as a decaying exponential, with diminishing reductions in BR to increasing $\Delta_{max}$. It can be observed that, depending on $n$, $\Delta_{max}$ = 64, 128, 256 are three good selections with balanced trade-offs.

$\delta_{max} = n$, the extra logic cells scale as $LC_{SH-EED} \propto log(n)$. The ROM occupation is the same as SH-EED case. As for the power consumption, we can only test the channel number under 1000 limited by the available onboard block RAMs. We have observed a more-than-linear growing extra power trend with increasing channel counts, comparing to the SH-DED. It needs less than 1% extra power at 10 channels, while that becomes 10% at 1000 channels.

In the case of limited max channel difference, tiny number of extra logic cells (No more than 10 logic cells) is need comparing to the $\Delta_{max} = n$ case. Its power upscaling also shares similar trending. The usage of ROM however can be highly constrained, which is translated to the reduced area occupation in future ASIC design. The power consumption is also reduced because of the reduced ROM usage. However, there is no clear clue on how the number of extra logic cells and power consumption are varying with $\Delta_{max}$ the power consumption is overall consistent. Therefore, one should trade off their ROM/area availably and BR requirement when selecting $\delta_{max}$. As the growing $\Delta_{max}$ has opposite effect on the ROM usage and BR reduction.

### 5 BR plots

For each considered number of channels, a range of $S$ values, $hs = 0, 6$ and the fixed $\Delta_{max}$ values, the BRs for all of the compression methods and implementations are given in Fig. 4 to 9 for BP values of 1 to 100 ms.

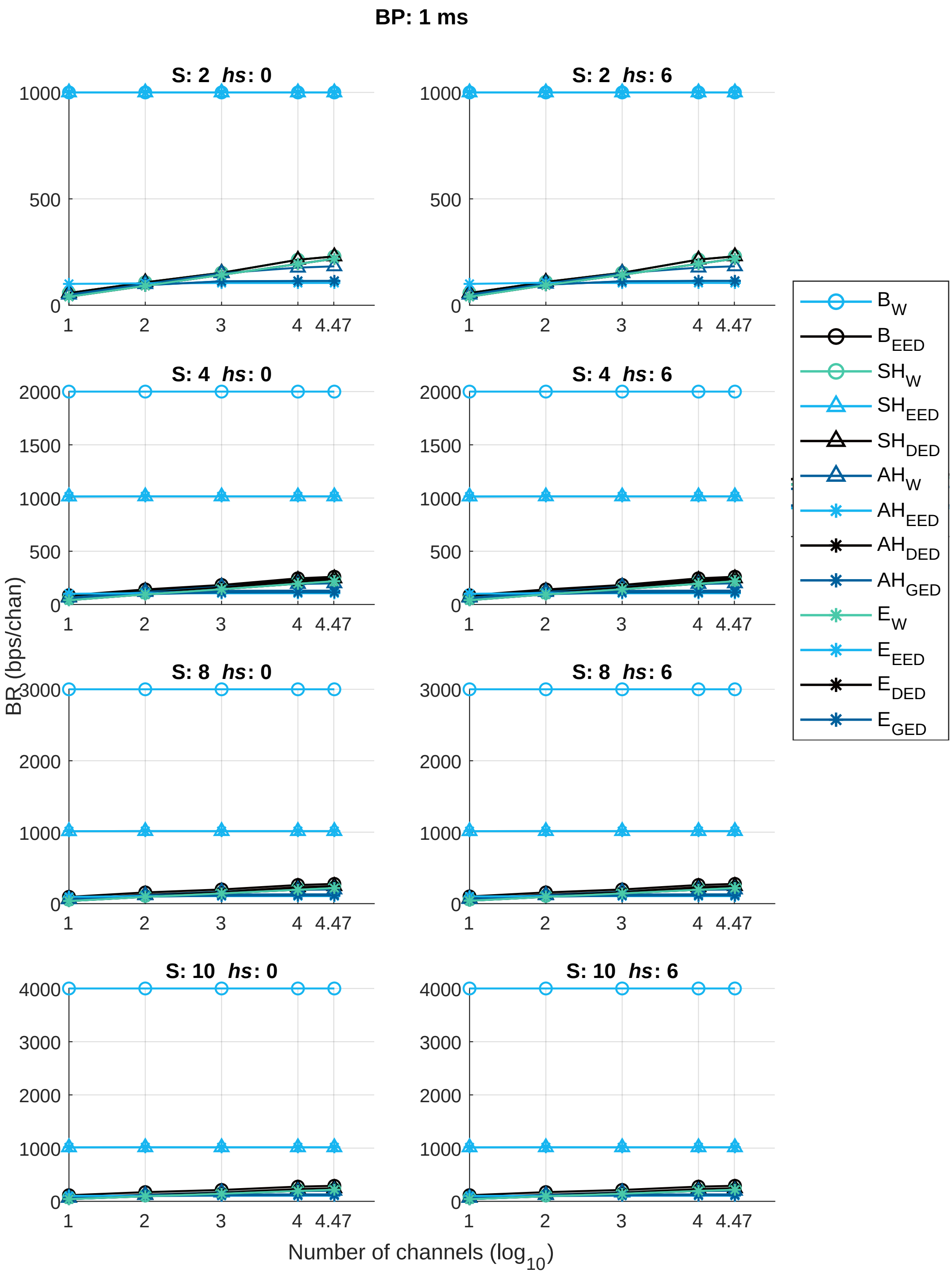


Figure 4: Bit Rates for communication schemes at a BP of 1 ms; $S \in [2,\ 4,\ 8,\ 10]$, $\Delta_{max} = min(n, 64)$.

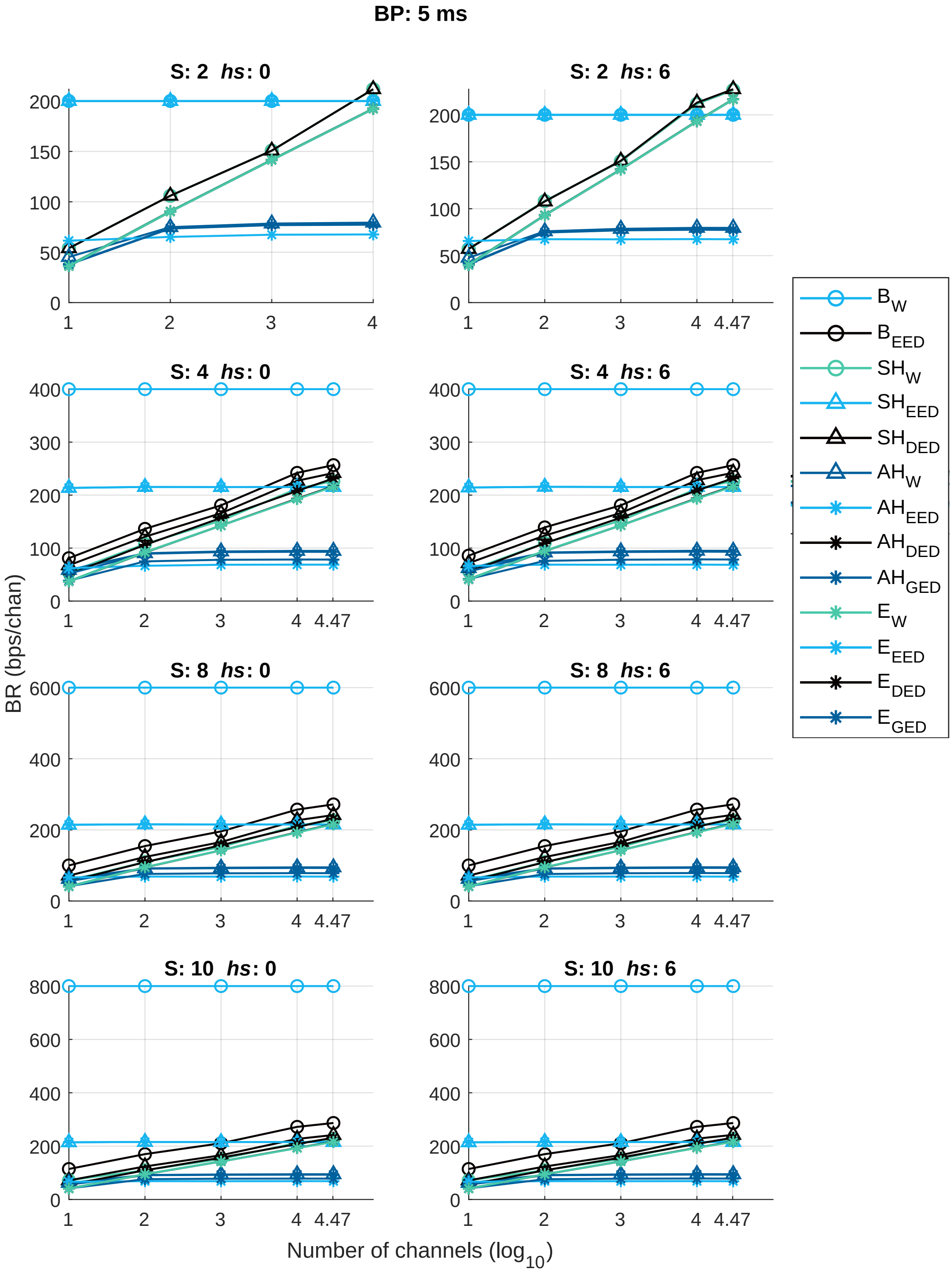


Figure 5: Bit Rates for communication schemes at a BP of 5 ms; $S \in [2,\ 4,\ 8,\ 10]$, $\Delta_{max} = min(n, 64)$.

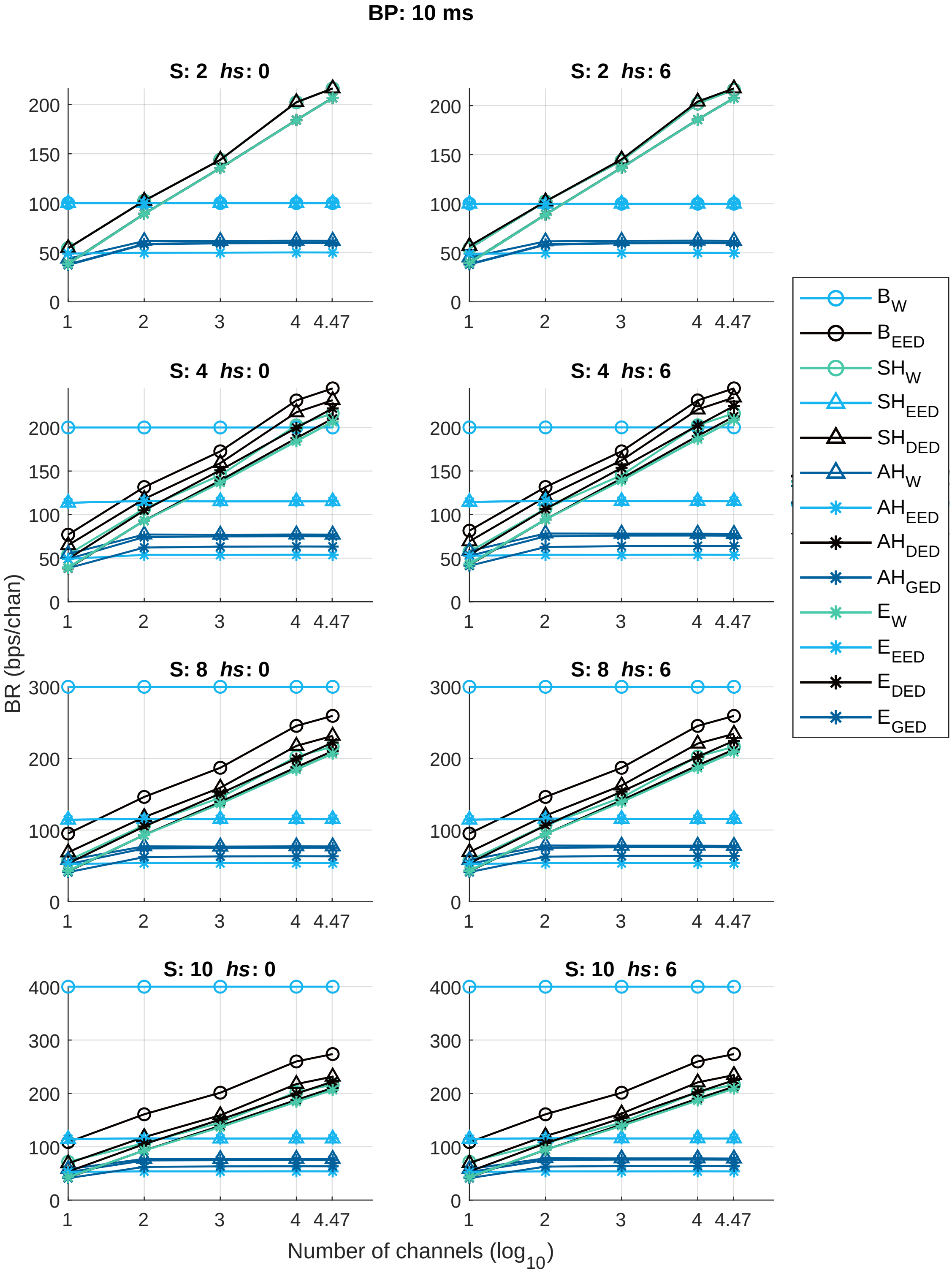


Figure 6: Bit Rates for communication schemes at a BP of 10 ms; $S \in [2,\ 4,\ 8,\ 10]$, $\Delta_{max} = min(n, 64)$.

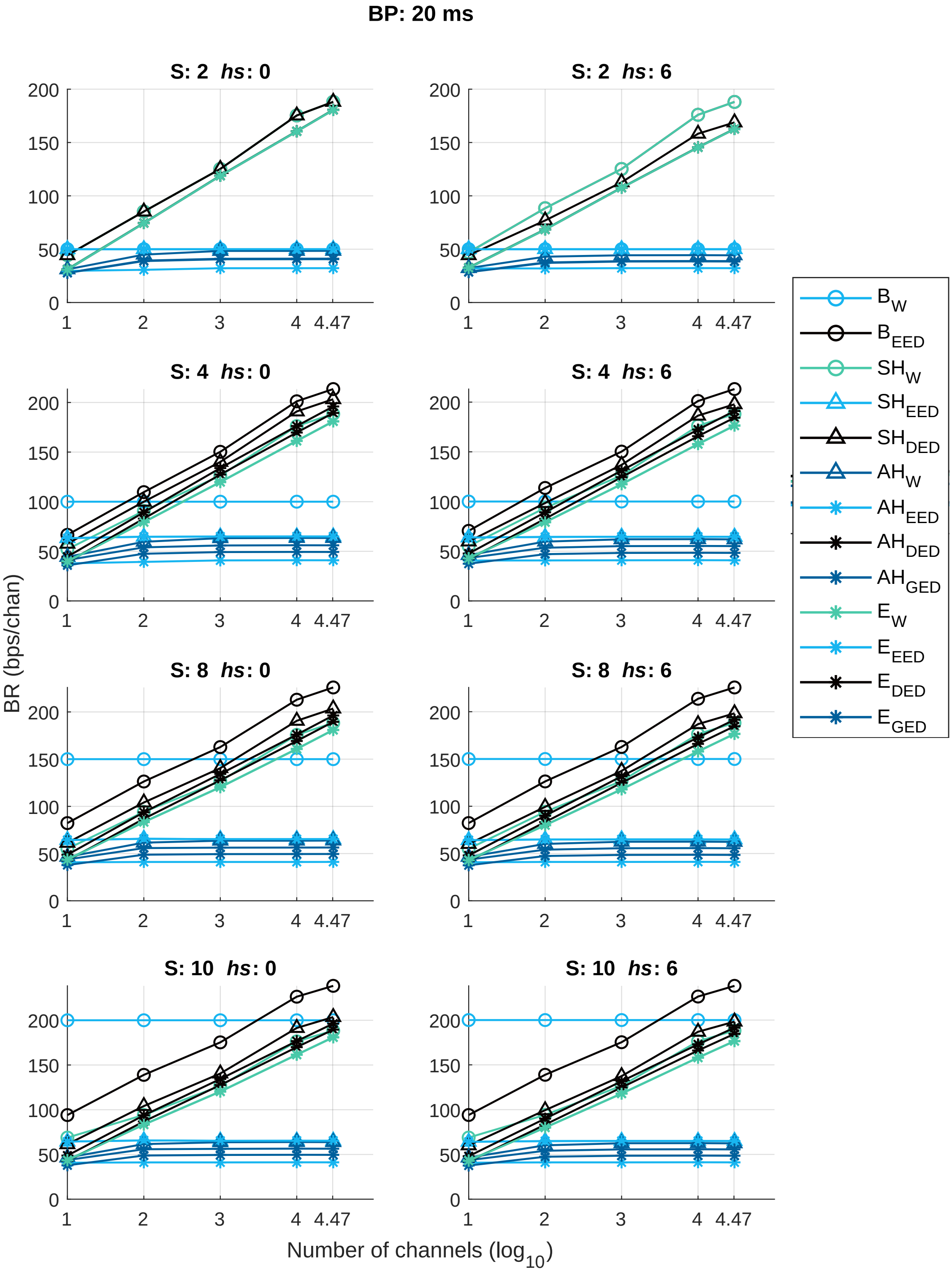


Figure 7: Bit Rates for communication schemes at a BP of 20 ms; $S \in [2, 4, 8, 10]$, $\Delta_{max} = min(n, 64)$.

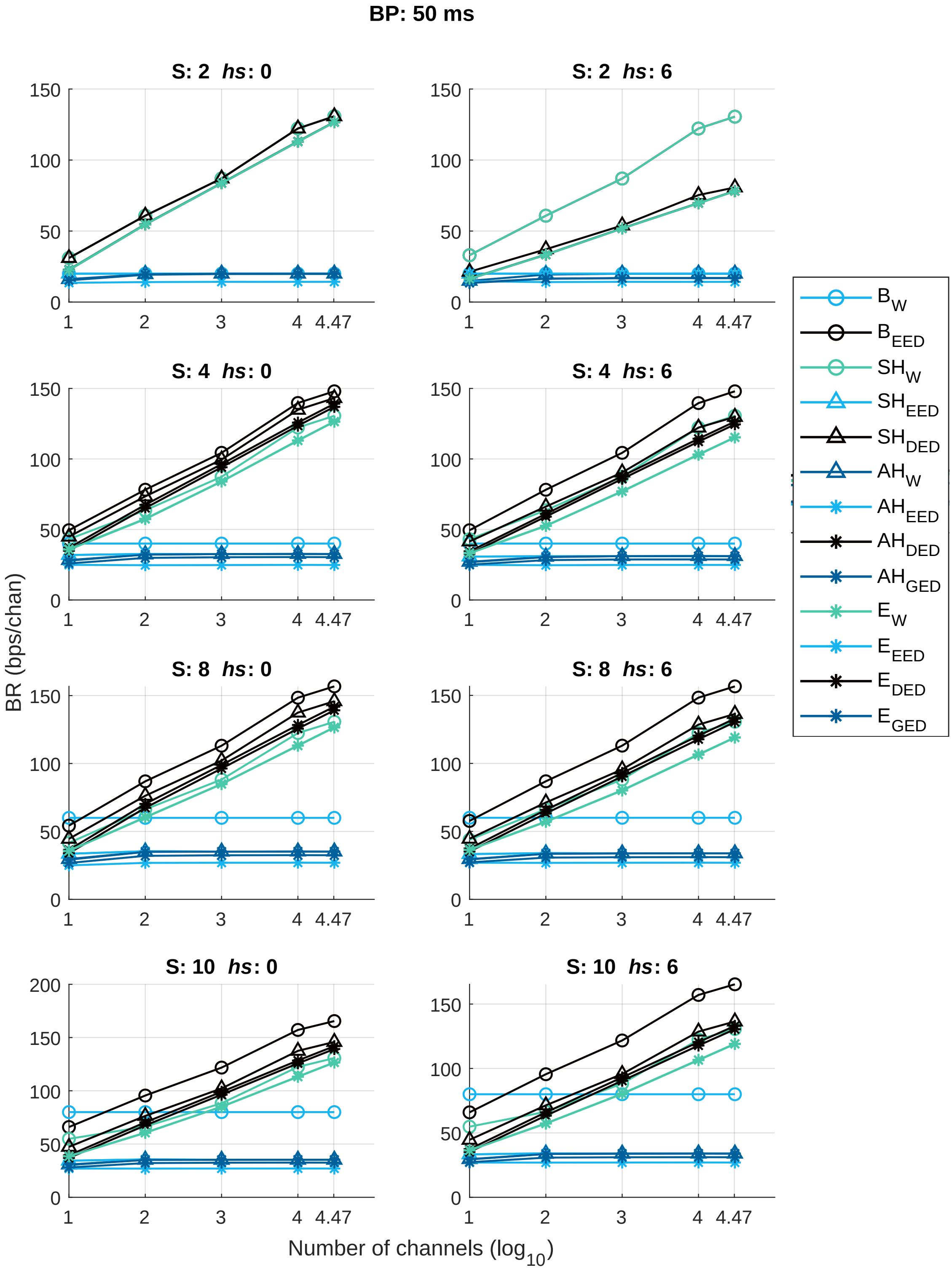


Figure 8: Bit Rates for communication schemes at a BP of 50 ms; $S \in [2,\ 4,\ 8,\ 10]$, $\Delta_{max} = min(n, 64)$.

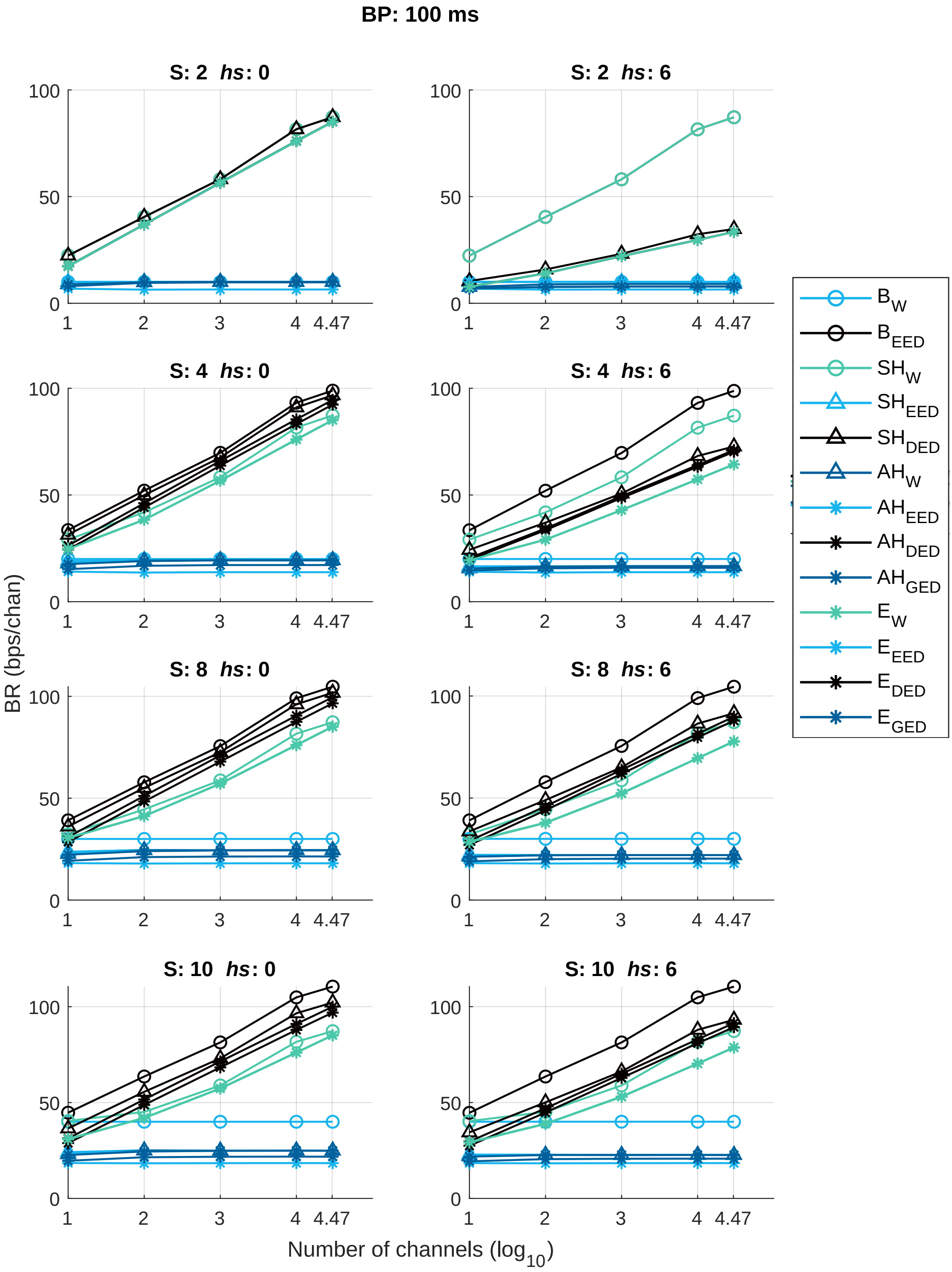


Figure 9: Bit Rates for communication schemes at a BP of 100 ms; $S \in [2,\ 4,\ 8,\ 10]$, $\Delta_{max} = min(n, 64)$.
